# Quantitation analysis by flow cytometry shows that *Wt1* is required for development of the proepicardium and epicardium

**DOI:** 10.1101/2020.10.06.329151

**Authors:** Christine Biben, Bette Borobokas, Mary Kamala Menon, Lynne Hartley, Richard Paul Harvey, Samir Taoudi, Owen William John Prall

## Abstract

The epicardium is a cell layer found on the external surface of the heart. During development it has an epithelial identity and contains progenitor cells for coronary smooth muscle and cardiac fibroblasts. The epicardium has been suggested to have therapeutic potential in cardiac repair. Study of epicardial development has been difficult because it is dynamic and morphologically complex. We developed a flow cytometry-based method to quantify cardiac development including the epicardial lineage. This provided accurate and sensitive analysis of (1) the emergence of epicardial progenitors within the proepicardium (2) their transfer to the heart to form the epicardium, and (3) their epithelial-to-mesenchymal transition (EMT) to create the subepicardium. Platelet-derived growth factor alpha (Pdgfra) and Wilms tumor protein (Wt1) have both been reported to be pro-mesenchymal during epicardial EMT. Quantitative analysis with flow cytometry confirmed a pro-mesenchymal role for Pdgfra but not for Wt1. Analysis of *Wt1* null embryos showed that they had (1) poor formation of proepicardial villi, (2) reduced transfer of proepicardial cells to the heart, (3) a discontinuous epicardium with poor epithelial identity, and (4) a proportionally excessive number of mesenchymal-like cells. This data shows that Wt1 is essential for epicardial formation and maintenance rather than being pro-mesenchymal.

## INTRODUCTION

Epicardial-derived cells have shown potential therapeutic role in adults for cardiac repair, either as progenitor cells for several cardiac cell types, or through paracrine mechanisms (Smart and Riley, 2012; Wang et al., 2015). This has generated interest in the mechanisms that guide their emergence and differentiation. The developmental origin of the epicardium (EPI) has been traced back in embryos to the proepicardium (PE), a transient structure lying caudal to the heart (Mikawa and Gourdie, 1996; Viragh and Challice, 1981). PE cells form villi that extend towards the heart, leading to cell transfer onto the myocardial surface via a cellular bridge (Rodgers et al., 2008) and/or release of clusters into the pericardial cavity (Komiyama et al., 1987; Sengbusch et al., 2002). These cells initially cover the myocardium as a monolayer, forming the early EPI. Some epicardial cells then leave this epithelial environment and adopt a mesenchymal fate, i.e. an epithelial-to-mesenchymal transition (EMT). This mesenchyme initially accumulates underneath the EPI (subepicardial mesenchyme or SEM) prior to invading the myocardium and differentiating into coronary smooth muscle cells and cardiac fibroblasts (Cai et al., 2008; Dettman et al., 1998; Zhou et al., 2008).

The transitions from PE to EPI, and from EPI to SEM, are highly dynamic and involve complex 3-dimensional changes. In the mouse this occurs over a 2-3 day period. Markers of the PE, EPI and SEM are limited and by themselves non-specific, at least initially. Some steps, like the accumulation of the SEM under the epicardium, show variability in timing and extent depending upon the location within the heart. For these reasons analysis of epicardial development in wild-type and mutant animals by 2-dimensional sections is qualitative and subject to significant sample error. We therefore screened potential cell surface antibodies for expression in PE, EPI and SEM. Using these markers and flow cytometry we developed a highly sensitive and quantitative method to analyze early epicardial development. Our results with this technique confirm and extend the known role of platelet-derived growth factor alpha (*Pdgfra*) in epicardial development, and identifies multiple novel roles for Wilms tumour 1 (*Wt1*).

## RESULTS

We searched for combinations of cell surface markers that would allow us to follow epicardial development by flow cytometry in embryos of any genotype. Initially, we used antibodies to the transcription factor WT1 and the *Wt1*-GFP transgene (*Wt1^GFPCre^* mice (Zhou et al., 2008)) to identify cells of the epicardial lineage during early cardiac development.

### Marker expression in the proepicardium (PE)

At embryonic day 9.5 (E9.5) in the mouse, epicardial progenitors reside in the PE. PE cells were identified at the ventral surface of the septum transversum by expression of WT1 (Fig. 1A). Integrin alpha4 (*Itga4*), has a critical role in early epicardium (Sengbusch et al., 2002), and was also expressed in PE cells (Fig. 1B). High levels of ITGA4 and WT1 were confined to the surface of the septum transversum, where PE villi are located. Unfortunately, neither of two anti-WT1 antibodies we used gave reliable signals by flow cytometry. Mice heterozygous for the *Wt1*^GFPCre^ allele (Zhou et al., 2008) were therefore used to identify WT1-expressing cells in the PE with this technique (Fig. 1C). We found that most *Wt1*-GFP^+^ cells were ITGA4^high^ and that all ITGA4^high^ cells were *Wt1*-GFP^+^, consistent with co-expression of WT1 and ITGA4 in PE villi. Quantification of either ITGA4^high^ PE cells by flow cytometry or manual counting of PE villi cells on H&E sections gave similar results (Fig. 1D). Epicardial progenitors begun to accumulate in the PE from 16-17 somite pairs and peaked around 24-28 somite pairs. PE cells also expressed PODOPLANIN (a known epicardial marker (Mahtab et al., 2008)), PDGFRA (Chong et al., 2011) and ALCAM (also found throughout the septum transversum (Asahina et al., 2011)) (Fig. S1A-C). In contrast, PDGFRB was not detected (Fig. S1D).

**Figure 1.**
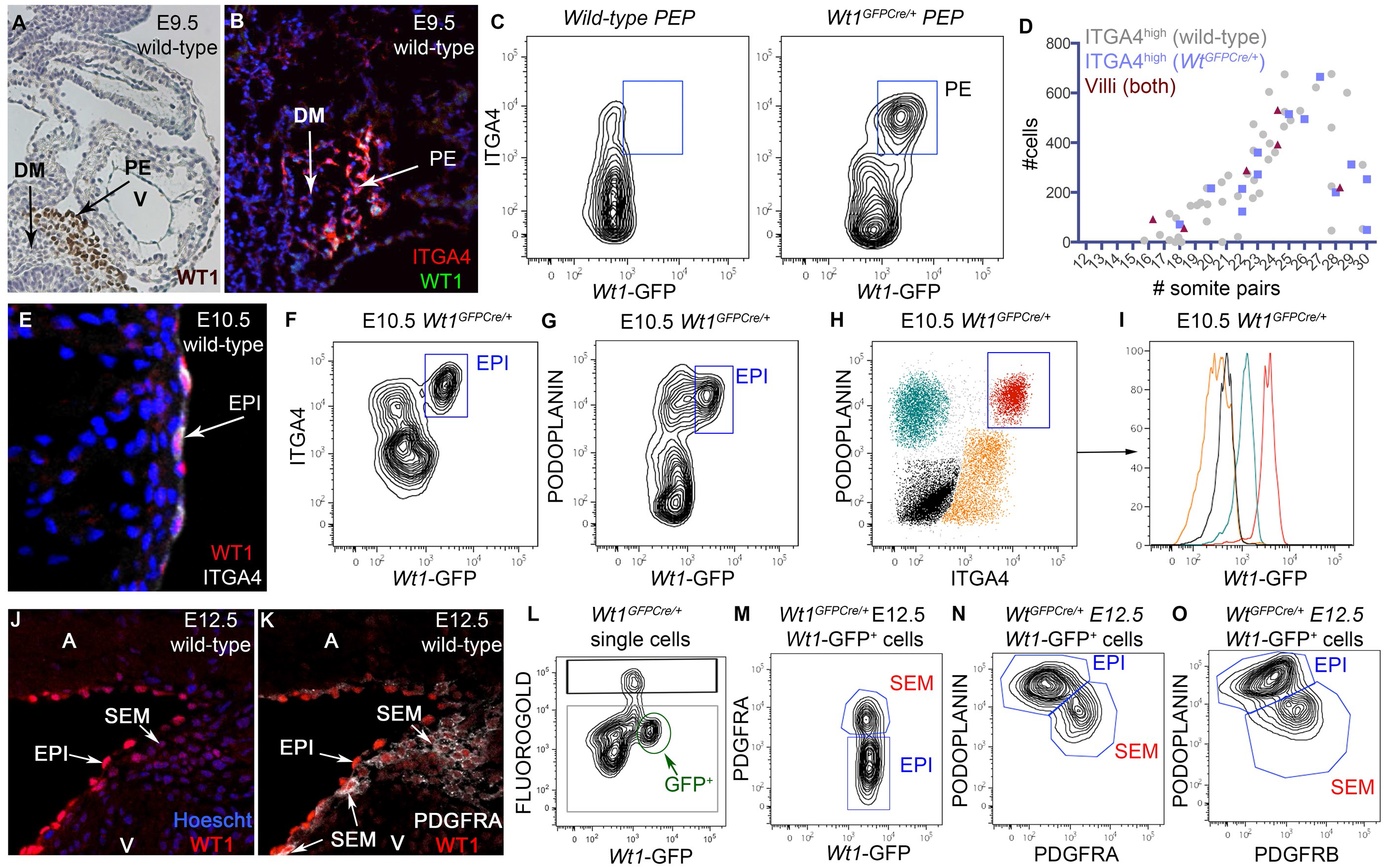
Quantitative analysis of early epicardial development. **A**. WT1 immunochemistry on E9.5 sections of wild-type embryo. High level of WT1 staining is found on PE villi while the dorsal mesenchyme (DM) shows only limited staining. V: ventricle. **B**. Immunofluorescence on E9.5 cryostat sections of wild-type embryo. High levels of ITGA4 and WT1 on PE villi. **C**. Flow cytometry analysis of wild-type and *Wt1^GFPCre/+^* PE preparations (PEP). ITGA4^high^ cells are Wt1-GFP^+^. **D**. Time course of proepicardial development: quantification of ITGA4^high^ cells by flow cytometry and counting of cells engaged in PE villi on H&E sections gives similar results. Genotypes are indicated. **E**. Immunofluorescence on E10.5 cryostat sections of wild-type embryo. High levels of ITGA4 and WT1 in EPI cells. **F-I**. Flow cytometry analysis of E10.5 *Wt1^GFPCre/+^* hearts. ITGA4^+^ *Wt1*-GFP^+^ EPI cells form a distinctive population (**F**). EPI cells are PODOPLANIN^+^ (**G**). PODOPLANIN vs ITGA4 identifies 4 populations: PODOPLANIN ^+^ ITGA4^+^ cells are the only *Wt1*-GFP^+^ cells (**H,I**). **J-K**. Immunofluorescence on E12.5 cryostat sections of wild-type heart. EPI cells are WT1^high^ PDGFRA^low^ and SEM cells are WT1^low^ PDGFRA^high^. A: atrium; V: ventricle. **L-O**. Flow cytometry analysis of E12.5 *Wt1^GFPCre/+^* hearts. (**L**) GFP^+^ cells can be gated specifically (green gate) out of live (grey) single cells. Black gate = dead cells. (**M-O**) Gating on *Wt1*-GFP^+^ cells only: two distinct populations are present: PDGFRA^+^ cells (red, SEM) are PODOPLANIN^low^ and PDGFRB^+^, PDGFRA^−^ cells (blue, EPI) are PODOPLANIN^high^ PDGFRB^−^.

### Marker expression in the pre-EMT (E10.5) Epicardium

WT1^+^ ITGA4^+^ cells were found on the myocardial surface from E9.5, rapidly covering the heart to create a continuous epicardial monolayer by E10.5 (Fig. 1E). As expected, *Wt1*-GFP^+^ ITGA4^+^ cells were detected in whole E10.5 *Wt1*^GFPCre/+^ hearts by flow cytometry (Fig. 1F). Since *Wt1*-GFP^−^ ITGA4^+^ cells were also present in the heart (Fig. 1F), additional markers were necessary to identify EPI in absence of the *Wt1*-GFPCre transgene. Early epicardial cells were found to retain expression of PODOPLANIN (Fig. 1G) and ALCAM (Fig. S1E), and the combination of either marker with ITGA4 was unique to the EPI (Fig. 1H,I, Fig. S1F,G). In addition, EPI cells were the only ITGA4^+^ PECAM^−^ cells in the heart at this stage (Fig. S1H).

### Marker expression in EPI and SEM during EMT

EMT of the EPI results in accumulation of the SEM. The SEM is initially most prominent in the atrioventricular and interventricular grooves (AVG, IVG). We found that the SEM expressed low levels of WT1 and high levels of PDGFRA, compared to the EPI (Fig. 1J,K). Flow cytometry showed that *Wt1*-GFP^+^ cells could be subdivided into PDGFRA^low/−^ and PDGFRA^high^ cells, putative EPI and SEM respectively (Fig. 1L,M). Mean *Wt1*-GFP epifluorescence was only slightly lower in SEM compared to EPI, suggesting that expression of GFP decreased more slowly than that of WT1. To confirm our identification of PDGFRA^low/−^ and PDGFRA^high^ cells as EPI and SEM respectively, we used the *Pdgfra*-GFP allele, which was compatible with intracellular flow cytometry, allowing us to look at keratin expression, a hallmark of epithelial (epicardium) tissues. PDGFRA is expressed in PE (Fig. S1B), then downregulated as cells form the EPI and upregulated in nascent SEM (Fig. 1J,K). Given that the *Pdgfra*-GFP transgene (and *Pdgfra* itself) is also expressed in valve mesenchyme and neural crest cells, unlike *Wt1*-GFP, ventricle preparations were used to ensure that all *Pdgfra*-GFP^+^ cells observed were of epicardial origin. Accumulation of SEM was lower than that observed with *Wt1*-GFP, as the AVG was not represented in ventricle fractions. Two *Pdgfra*-GFP^+^ populations could be recognized from E11.5, a *Pdgfra*-GFP^low^ PDGFRA^low/−^ (blue arrow, Fig. S2A) and a *Pdgfra*-GFP^high^ PDGFRA^+^ (red gate, Fig. S2A) population. Expression of cytokeratins KRT8 and KRT19 confirmed *Pdgfra*-GFP^low^ PDGFRA^low/−^ population as the epicardium (EPI) (Fig. S2B,C), while expression of mesenchymal markers (PDGFRA, PDGFRB) was confined to the *Pdgfra*-GFP^high^ PDGFRA^+^ KRT8^−^ KRT19^−^ population (SEM) (Fig. S2A-D). Differential expression of PODOPLANIN (high in EPI) and ITGA4 (high in SEM) was also observed (Fig. S2E,F).

Differential expression of PDGFRB, ITGA4 and PODOPLANIN between EPI and SEM was also observed using *Wt1*-GFP (Fig. S1I-K). Combinations of PODOPLANIN with PDGFRA, PDGFRB or ITGA4 were all sufficiently sensitive to separate EPI and SEM within the *Wt1*-GFP^+^ population (Fig. 1N,O, Fig. S1L).

### Monitoring epicardial EMT in wild-type embryos

Combinations of PODOPLANIN, PDGFRA, ITGA4 and PDGFRB were able to uniquely identify EPI and SEM in presence of either the *Wt1*-GFP or *Pdgfra*-GFP transgenes. To identify strategies that would allow us to do this without relying on any transgene, we profiled the expression of those markers in various cardiac derivatives.

### Identification of non-epicardial cells in the embryonic heart

Transgenic lines that labeled endothelium and endothelial derivatives such as valve mesenchyme (*Tie2*Cre (Kisanuki et al., 2001)), blood (*Tie2*Cre (Kisanuki et al., 2001)), epicardial and valve mesenchyme (*Pdgfra*-GFP (Hamilton et al., 2003)), myocardium (*Nkx2-5*Cre (Stanley et al., 2002)) and neural crest derivatives (*Wnt1*Cre (Chai et al., 2000)) were used to identify these lineages. Well described markers such as PECAM (endothelium), PDGFRA and PDGFRB (mesenchymal cells (Andrae et al., 2008)), Ter119 (erythrocytes (Chao et al., 2015)), ALCAM and VCAM (myocardium (Elliott et al., 2011; Kwee et al., 1995; Plavicki et al., 2014)), PODOPLANIN and ITGA4 (epicardium (Mahtab et al., 2008) (Sengbusch et al., 2002)) were included to identify combination of markers specific to each lineage. Dissociation of E10.5 hearts showed a cell population with high autofluorescence in the 405 and 488 channels that did not express epicardial (ITGA4, Fig. S3A), mesenchymal (PDGFRA, Fig. S3B) or endothelial (PECAM) markers, but was 100% recombined in *Nkx2-5Cre/+ ZRed/+* hearts (Fig. S3C). This population also expressed VCAM and ALCAM (Fig. S3D,E) consistent with early myocardium. Early myocardium also expressed PODOPLANIN (Fig. S3F). In *Tie2Cre/+ RosaYFP/+* hearts there were two populations of YFP-expressing cells. *Tie2*Cre-YFP^mid^ Ter119^+^ cells corresponded to erythrocytes (Fig. S3G,H). *Tie2*Cre-YFP^high^ cells were further subdivided by PECAM expression into three sub-populations. Endothelium was PECAM^high^ ITGA4^low/−^ PDGFRA^−^ *Wt1*-GFP^−^, (Fig. S3G-K blue). Endothelial-derived valve mesenchyme was PECAM^low^ ITGA4^mid^ PDGFRA^high^ *Wt1*-GFP^−^ (Fig. S3G-K orange). The third group was PECAM^−^ ITGA4^high^ PDGFRA^+^ ALCAM^+^ (Fig. S3G-L green), as bona fide EPI, and most likely corresponded to the rare EPI cells deleted in the PE in *Tie2Cre/+ Rosa26R/+* embryos (Fig. S3M). Neural crest cells were identified as *Wnt1*Cre-YFP^+^ and expressed mesenchymal markers including PDGFRA and ITGA4, but were low in ALCAM (Fig. S3N).

In addition to epicardial development, the PDGFRA/ITGA4/ALCAM or PODOPLANIN/PECAM combination allowed the monitoring of other cardiac events such as valve EMT. Removal of the outflow tract (OFT) and its corresponding cushions permitted quantification of atrioventricular (AV) cushions EMT in the remaining fraction, with no contamination from either OFT mesenchyme or neural crest cells (Fig. S4A,B). AV valve mesenchyme (PECAM^low^ ITGA4^mid^) accumulated rapidly between 22-32 somite pairs (Fig. S4C). OFT cushions EMT could also be followed accurately in OFT preparations by using *Wnt1*Cre to separate neural crest from the valve mesenchyme (Fig. S4D).

### Quantification of epicardial development in wild-type samples

Some neural crest, and to a lesser degree valve mesenchyme, started to contaminate the EPI gate at E11.5 (Fig. S5A-E). Ventricle preparations that exclude all valves and the OFT were used from this stage onwards as in these the number of either cell type was negligible (Fig. S5F-J). Up to EMT, ALCAM and PODOPLANIN had been interchangeable for identifying the epicardial lineage as a whole, however the differential PODOPLANIN expression in EPI and SEM made it more attractive than ALCAM (whose levels were less affected, see below) from E11.5. EPI and SEM were therefore quantified in wild-type ventricles using the following strategy. PODOPLANIN^+^ ITGA4^+^ cells were gated out of single viable cells (Fig. 2A). Contaminating myocardial cells were then removed using their autofluorescence in the 488 channel (Fig. 2B). The remaining cells were then of the epicardial lineage, and relative levels of PODOPLANIN and PDGFRB were used to separate EPI from SEM (Fig. 2C). In *Wt1*-GFPCre^+^ samples, GFP^+^ cells were selected first on a fluorogold vs GFP plot (Fig. S5K-M) and then subjected to the strategy described in Fig. 2A&C.

**Figure 2.**
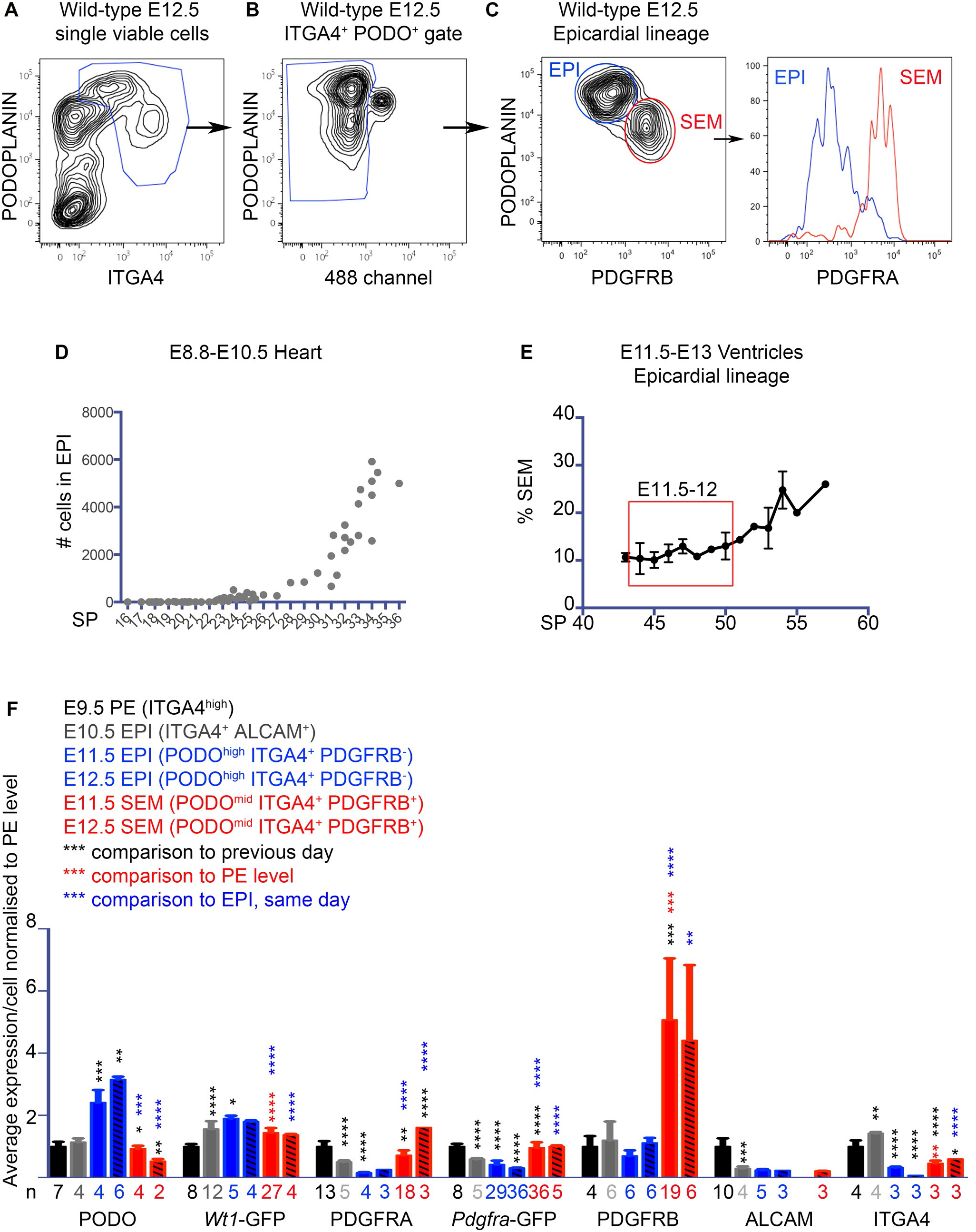
Endogenous and reporter marker expression during early epicardial development. **A-C**. Gating strategy for epicardial derivatives in wild-type E11.5-E12.5 ventricles. PODOPLANIN^+^ ITGA4^+^ cells are gated of single viable cells (**A**). PODOPLANIN vs GFP (488) channel allows gating out of myocardial cells (autofluorescent) (**B**). PODOPLANIN vs PDGFRB allows separation of PODOPLANIN^high^ PDGFRB^−^ EPI cells (blue) and PODOPLANIN^mid^ PDGFRB^+^ SEM cells (red) (**C**). Differential expression of PDGFRA (not used for gating) confirms EPI and SEM identities. **D**. Accumulation of epicardial cells (ITGA4^+^ ALCAM^+^) with time in the heart. SP: somite pairs. Each dot represents an embryo. **E**. Proportion of SEM cells (PODO^mid^ ITGA4^+^ PDGFRB^+^) within the epicardial lineage (PODO^+^ ITGA4^+^) between E11.0 and E13. SP: somite pairs. **F**. Mean marker expression per cell by flow cytometry in the PE, EPI and SEM. All markers normalised to a relative expression of 1 in the PE. Gating strategies and stages are indicated. PODO = PODOPLANIN. Black = significance compared to the previous day (E9.5 PE -> E10.5 EPI -> E11.5 EPI + E11.5 SEM). For E12.5 EPI and SEM samples, the previous day samples are E11.5 EPI and SEM respectively. Red = significance compared to the PE level. Blue = significance compared to the EPI of the same day. **E,F**: Mean ± SD are displayed. Statistics: Student’s t-test. * p<0.05, ** p <0.01, *** p <0.001, **** p<0.0001. n= number of independent samples.

Quantification of wild-type epicardial development showed that PE cells reached the heart from 22 somite pairs onwards (Fig. 2D). EPI cells expansion showed 2 phases: an initial slow phase concomitant with the cell transfer followed by a step of vigorous expansion. Ki67 labeling confirmed that EPI cells were undergoing rapid proliferation at that stage (85.3 ± 0.85 % of cells in G1 to M cell cycle phases, n=8). Accumulation of SEM in the ventricles was monitored from E11.0 (Fig. 2E) and showed a relatively stable ratio of EPI to SEM until E12.0, after which SEM expansion was accelerated, probably due a combination of EMT and mesenchymal proliferation.

Mean expression per cell was analyzed for various markers in the PE, EPI and SEM (Fig. 2F). Transition from the PE (E9.5) to the early (pre-EMT) EPI (E10.5) was accompanied by increased *Wt1*-GFP, and decreased PDGFRA, *Pdgfra*-GFP and ALCAM. This trend continued in the EPI from E11.5, accompanied by an increase in PODOPLANIN and a decrease in ITGA4. EMT of the EPI to create the SEM was associated with decreased PODOPLANIN and *Wt1*-GFP, and increased PDGFRA, *Pdgfra*-GFP, PDGFRB and ITGA4 expression.

These experiments when taken together show that epithelial (EPI) status correlated with high levels of WT1, *Wt1*-GFP and PODOPLANIN and low levels of PDGFRA, *Pdgfra*-GFP, PDGFRB and ITGA4. Mesenchymal (SEM) status correlated with the reciprocal expression profile. The PE showed expression of both epithelial (WT1, *Wt1*-GFP, PODOPLANIN) and mesenchymal (PDGFRA, *Pdgfra*-GFP and ITGA4, but not PDGFRB) markers, and uniquely expressed high levels of ALCAM.

### Quantification of epicardial development confirms that *Pdgfra* is required for development of the subepicardial mesenchyme

PDGF signaling has been described as critical for epicardial EMT (Martinez-Estrada et al., 2010; Smith et al., 2011). PDGFRA is expressed in PE and pre-EMT EPI, before being markedly increased in nascent SEM (Fig. S1C, 1J-K, S6A-C). We used two alleles of *Pdgfra* to investigate its role in epicardial development: *Pdgfra*^GFP^, a histone 2B-GFP knock-in null allele (Hamilton et al., 2003), and *Pdgfra*^*floxed*^, a hypomorphic allele (Tallquist and Soriano, 2003). The *Pdgfra*^*floxed*^ allele is also conditional-null following Cre-mediated recombination. We generated three mutant models: *Pdgfra*^*GFP/GFP*^ null embryos, *Pdgfra*^*GFP/floxed*^ hypomorphic embryos and so called “epicardial-deleted” mutants, *Gata5Cre/+ Pdgfra^floxed/floxed^*. *Pdgfra^GFP/+^* heterozygous mice developed without a gross morphological phenotype as reported (Hamilton et al., 2003). *Pdgfra*^*GFP/floxed*^ hypomorphic embryos were obtained at Mendelian ratios at E12.5. *Pdgfra*^*GFP/GFP*^ null embryos had a more severe phenotype, with up to 75% mortality by E10.5, as previously described (Soriano, 1997). Rare “healthy” null embryos could still be obtained at E12.5, before developing fatal cranial hemorrhages at E14. Although a SEM began to accumulate at E11.5 in *Pdgfra*^*GFP/GFP*^ null hearts (Fig.3A,B), SEM was reduced at E12.5 in both hypomorphic and null embryos (2.1 and 2.4-fold less than control heterozygous mice, respectively, p<0.001, Fig. 3C). The average PDGFRA expression/cell in *Pdgfra^floxed/+^, Pdgfra^GFP/+^, Pdgfra^floxed/floxed^* and *Pdgfra*^*GFP/floxed*^ embryos was 69%, 36%, 34% and 20% of wild-type respectively (after subtracting the *Pdgfra*^*GFP/GFP*^ background, Fig. 3D), confirming that the floxed allele was hypomorphic, and that *Pdgfra*^*GFP/floxed*^ embryos expressed very little PDGFRA. We did not find evidence of compensatory mechanisms via upregulation of *Pdgfra* or *Pdgfrb*: the level of GFP fluorescence in *Pdgfra*^*GFP/floxed*^ embryos was statistically undistinguishable from that of *Pdgfra^GFP/+^* embryos (*Pdgfra^GFP/+^* n=10, *Pdgfra*^*GFP/floxed*^ n=6, p=0.38) and *Pdgfra*^*GFP/GFP*^ embryos averaged 1.9-fold that level (*Pdgfra^GFP/+^* n=13, *Pdgfra*^*GFP/GFP*^ n=5, p<0.0001). In addition PDGFRB levels were unchanged (*Pdgfra^GFP/+^* n=10, *Pdgfra*^*GFP/floxed*^ n=6, *Pdgfra*^*GFP/GFP*^ n=5, p=0.50-0.79). Loss of SEM did not appear to be a consequence of excessive apoptosis (Fig. 3E). However, SEM proliferation was mildly reduced compared to that of heterozygous embryos (Fig. 3F). Proliferation of the EPI, which expresses very little PDGFRA at this stage, was unaffected (Fig. 3F).

**Figure 3.**
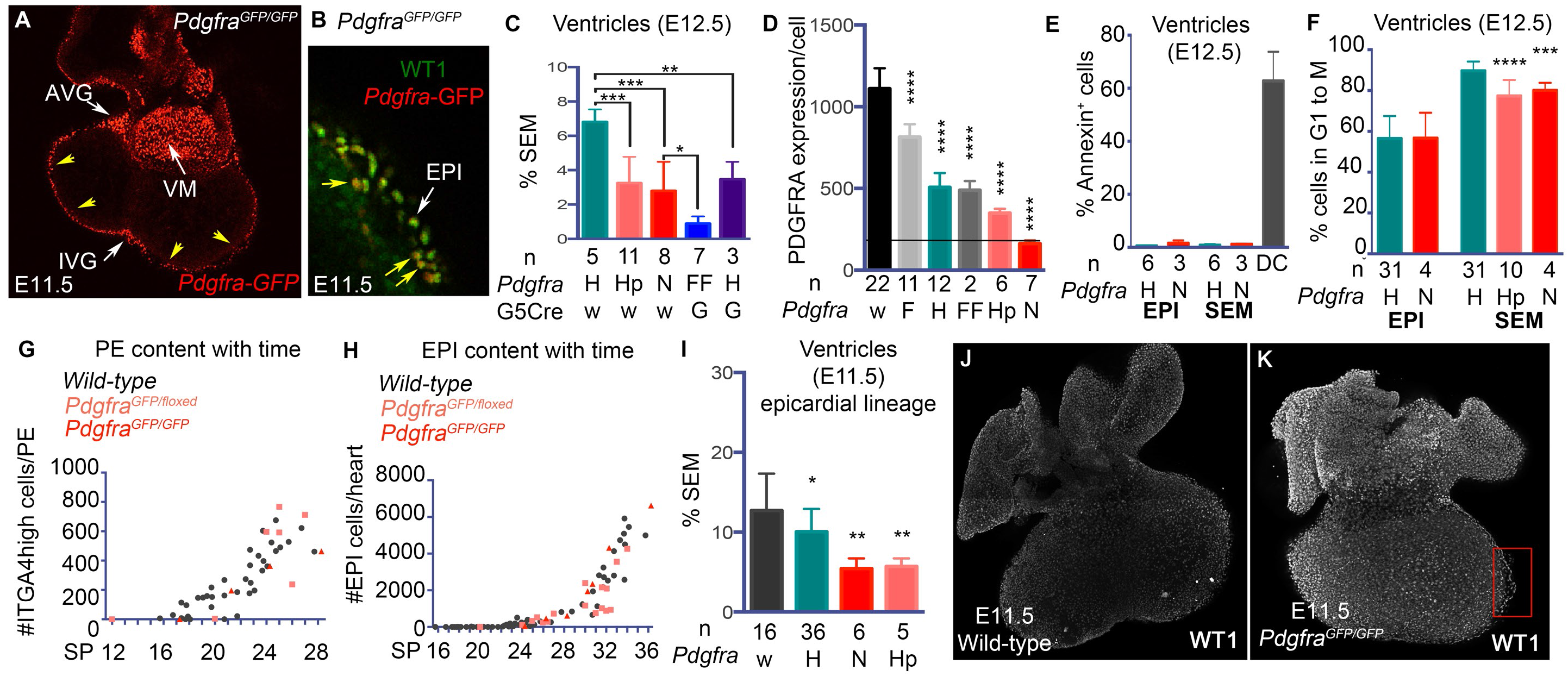
Embryos lacking Pdgfra have a mild reduction in SEM. **A,B**. Optical sections through a whole mount *Pdgfra*^*GFP/GFP*^ E11.5 heart stained with WT1. Only GFP channel is displayed in **A**. Presence of SEM (*Pdgfra*-GFP^high^ cells) is obvious at the level of the atrioventricular groove (AVG) and interventricular groove (IVG), as well as underneath the ventricular epicardium (**B**). NB: *Pdgfra*-GFP is also expressed in valve mesenchyme (VM). **C**. %SEM in E12.5 ventricles. Mean ± SD are displayed. Statistics: Student’s t-test. * p<0.05, ** p <0.01, *** p <0.001. n= number of independent samples. Genotypes are indicated (H: heterozygote; Hp: hypomorph; N: null; FF: floxed/floxed; G: *Gata5*Cre). **D**. PDGFRA expression per cell (arbitrary units) in various genotypes. Genotypes: w = wild-type, F = *Pdgfra^floxed/+^*, H = heterozygotes (*Pdgfra^GFP/+^*), FF = *Pdgfra^floxed/floxed^*, Hp = hypomorphs (*Pdgfra*^*GFP/floxed*^), N = nulls (*Pdgfra*^*GFP/GFP*^). Statistics: Student’s t-test, comparison to wild-type. **E**. Proportion of Annexin^+^ cells in E12.5 ventricles of *Pdgfra^GFP/+^* (H) and *Pdgfra*^*GFP/GFP*^ (N) embryos. DC: dead cells. **F**. Proportion of EPI and SEM cells actively in cell cycle (Ki67^+^) by flow cytometry at E12.5 in *Pdgfra^GFP/+^*(H), *Pdgfra*^*GFP/floxed*^ (Hp) and *Pdgfra*^*GFP/GFP*^ (N) embryos. Statistics: Student’s t-test, comparison to matching *Pdgfra^GFP/+^*(H) sample. **G,H**. Number of proepicardial (**G**, ITGA4^high^) and epicardial cells (**H**, ITGA4^+^ ALCAM^+^) by flow cytometry with time (somite pairs, SP) in wild-types, *Pdgfra*^*GFP/floxed*^ and *Pdgfra*^*GFP/GFP*^ embryos. Each dot represents an embryo. **I**. %SEM within the epicardial lineage (PODO^mid^ ITGA4^+^ PDGFRB^+^) of E11.5 ventricles. Statistics: Student’s t-test, comparison to wild-type. **J,K**. WT1 expression (white) by whole-mount immunofluorescence in wild-type (**J**) and *Pdgfra*^*GFP/GFP*^ null (**K**) hearts at E11.5. Red box = epicardial bubbles. **C-F,I**: Mean ± SD are displayed. Statistics (Student’s t-test): * p<0.05, ** p <0.01, *** p <0.001, **** p<0.0001. n= number of independent samples.

To assess the origin of this defect, early epicardial development was quantified in both *Pdgfra*^*GFP/GFP*^ and *Pdgfra*^*GFP/floxed*^ embryos. Embryos with morphological cardiac defects were excluded from this analysis. Accumulation of PE and early EPI was normal (Fig. 3G,H). SEM accumulation was reduced from its earliest point of accumulation at E11.5 (Fig. 3I), suggesting a defect in epicardial EMT. Flow cytometry showed normal expression of *Pdgfra*-GFP, PODOPLANIN, ITGA4 and PDGFRB in mutant EPI (Fig. S6D). However, 3D confocal analysis of whole-mount stained hearts revealed increased WT1 staining in *Pdgfra* mutant EPI at E11.5 (n=4/4, Fig. 3J,K). Morphological observation, as well as confocal imaging, showed that *Pdgfra*^*GFP/GFP*^ epicardium often appeared “bubbly” from E11.5 onwards (red box, Fig. 3K). Although this could be observed in control hearts as well as E11.5, this was more severe and persistent in the mutants. These experiments confirmed a role for *Pdgfra* in SEM formation, and suggest that this is at least in part due to decreased SEM proliferation and possibly also due to reduced epicardial EMT.

We attempted to delete *Pdgfra* specifically within the epicardial lineage with the *Gata5*Cre transgene. However these experiments were complicated by progressive transgene inactivation in stud males. Therefore efficiency of deletion was monitored with an anti-PDGFRA antibody in every sample, and only embryos with >90% deletion were analyzed. *Gata5Cre/+ Pdgfra^floxed/floxed^* embryos died at E14.5 with epicardial effusion and subepicardial blood pooling (n=3/3). At E12.5, *Gata5Cre/+ Pdgfra^floxed/floxed^* embryos showed no evidence of SEM, whether by flow cytometry (n=7/7, p<0.0001, Fig. 3C) or on sections (n=3/3, Fig. S6E,F), and had a stunted coronary endothelial network (n=3/3, Fig. S6G,H). This phenotype was significantly more severe than that of *Pdgfra* null embryos (Fig. 3F). In addition, *Pdgfra^GFP/+^ Gata5Cre/+* embryos had significantly less SEM than *Pdgfra^GFP/+^* embryos (p= 0.0015, Fig. 3C). This data showed that the *Gata5Cre* transgene itself interfered with epicardial development independently of deletion of a target floxed allele, and is therefore unsuitable for studying epicardial development.

### *Wt1* is required for development of the epicardium

*Wt1* is proposed to be required for development of the SEM (Martinez-Estrada et al., 2010; Moore et al., 1999; von Gise et al., 2011). *Wt1^GFPCre^* heterozygous null (von Gise et al., 2011) embryos (*Wt1^GFPCre/+^*) showed normal epicardial development (Fig. S7A-C). *Wt1^GFPCre/GFPCre^* homozygous null embryos became moribund from E13.5 as previously described (Moore et al., 1999). To avoid indirect effects they were not analyzed after E12.5. Whole mount confocal sections of E11.5 hearts labeled with an anti-GFP antibody showed a discontinuous epicardium in *Wt1^GFPCre/GFPCre^* embryos (Fig. 4A-C, n=4/4). Quantification of *Wt1*-GFP^+^ cells confirmed decreased EPI content in *Wt1^GFPCre/GFPCre^* hearts at E10.5 and E11.5 (Fig. 4D). Whole-mount staining and flow cytometry both showed that the discontinuous epicardial layer of E11.5 *Wt1^GFPCre/GFPCre^* ventricles contained cells with a broad range of GFP and PDGFRA levels and cell morphologies (Fig. 4E-H). PDGFRA^high^ Wt1-GFP^low^ cells showed multiple protrusions consistent with mesenchyme (red arrows, Fig. 4H). PDGFRA^low^ Wt1-GFP^high^ cells were flatter and rounder consistent with epithelial EPI cells (blue arrows, Fig. 4H). As the mesenchymal (PDGFRA^high^ Wt1-GFP^low^) cells were located on the surface of the myocardium, and not between EPI and myocardium as in controls, they were termed “SEM-like” cells. In *Wt1^GFPCre/GFPCre^* null mutants, SEM-like cells were increased in proportion within the epicardial lineage, whether in whole hearts, or ventricle preparations (Fig. 4I, n=11/11, p<0.0001).

**Figure 4.**
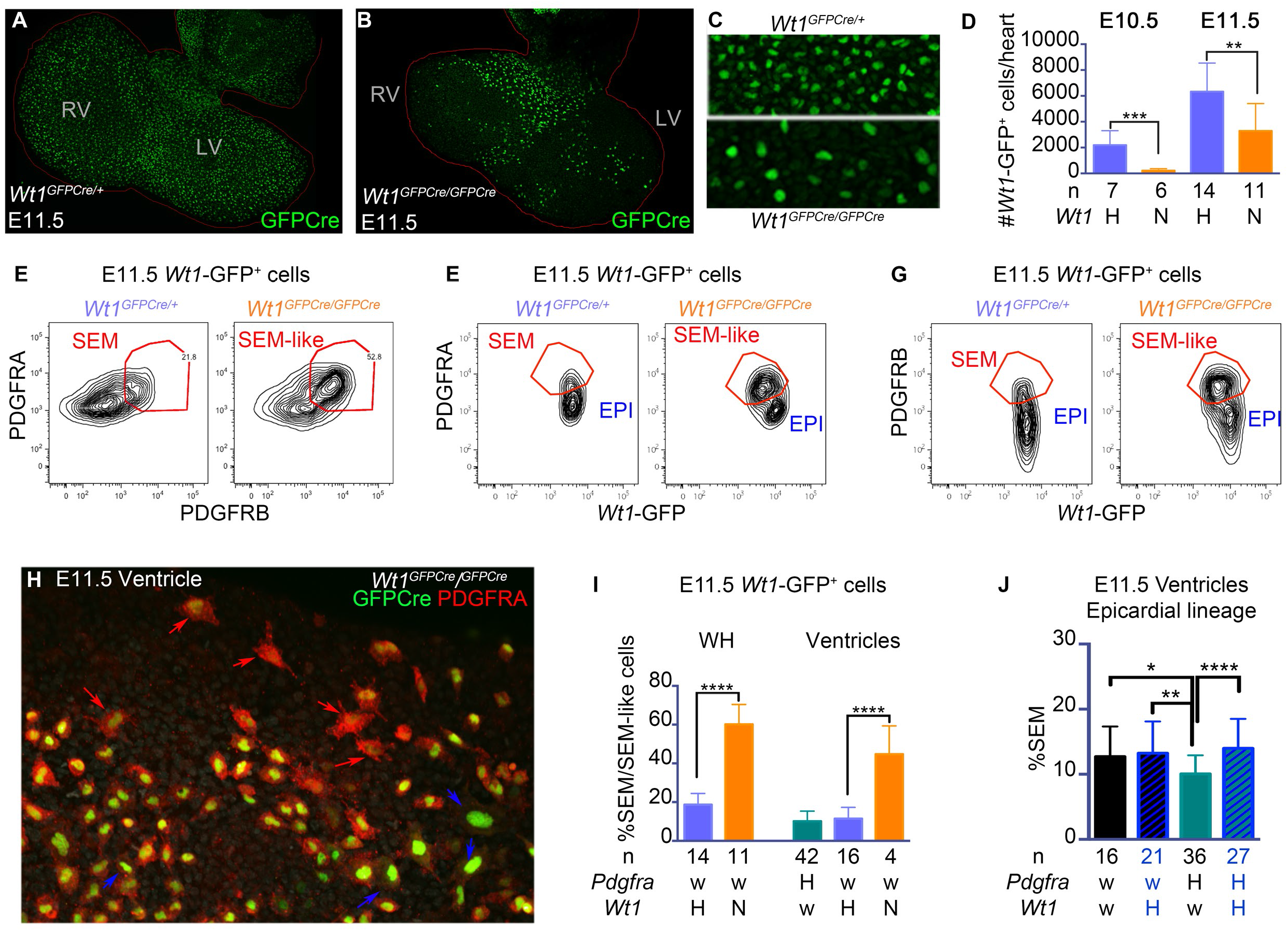
*Wt1* null embryos have a discontinuous epicardium and increased mesenchymal content. **A-C**. GFPCre protein expression (anti-GFP antibody) by whole-mount immunofluorescence in *Wt1^GFPCre/+^* (**A**, n=5) and *Wt1^GFPCre/GFPCre^* (**B**, n=4) hearts at E11.5. Red line = heart outline. (**C** = enlargement of **A** and **B**). **D**. Number of *Wt1*-GFP^+^ cells in *Wt1^GFPCre/+^* (H) and *Wt1^GFPCre/GFPCre^* (N) hearts at E10.5 and E11.5. **E**. PDGFRA and PDGFRB expression by flow cytometry in *Wt1*-GFP^+^ cells from *Wt1^GFPCre/+^* and *Wt1^GFPCre/GFPCre^* hearts at E11.5. **F,G**. PDGFRA, PDGFRB and *Wt1*-GFP expression in *Wt1^GFPCre/+^* and *Wt1^GFPCre/GFPCre^* hearts by flow cytometry at E11.5. **H**. PDGFRA and *Wt1*-GFP expression in *Wt1^GFPCre/GFPCre^* embryos by whole-mount immunofluorescence. PDGFRA^high^ *Wt1*-GFP^low^ SEM-like cells (red arrows) are mixed with PDGFRA^low^ *Wt1*-GFP^high^ EPI cells (blue arrows) on the surface of the heart. **I**. Proportion of SEM-like cells within *Wt1*-GFP^+^ cells at E11.5 by flow cytometry in whole heart (WH) and ventricle-only preparations. Genotypes are indicated. **J**. Proportion of SEM (PODOPLANIN^+^ ITGA4^+^ PDGFRB^+^) within the epicardial lineage (PODOPLANIN^+^ ITGA4^+^) of the ventricles at E11.5 (44-50sp). **I,J**: alleles for each gene: w = wild-type, H = heterozygote (either *Pdgfra^GFP/+^* or *Wt1*^*GFPCre/+*^) **D,I,J**: Mean ± SD are displayed. Statistics (Student’s t-test): * p<0.05, ** p <0.01, *** p <0.001, **** p<0.0001. n= number of independent samples.

### *Wt1* is required for epithelial identity of the epicardium

Decreased EPI and increased SEM-like cells in *Wt1* nulls suggested that WT1 has a pro-epithelial role, either by repressing epicardial EMT, or alternatively impairing the epithelial character of the EPI. Since *Pdgfra* null embryos showed increased WT1 staining and a reduced EMT, we investigated whether lowering WT1 expression in *Pdgfra* mutants could rescue the EMT defect. Ventricle samples from healthy embryos between 44-50 somite pairs were analyzed to ensure reliably measurable amounts of SEM and stage matching. Cells of the epicardial lineage were gated as described in Fig. 2 and the levels of PODOPLANIN vs PDGFRB used to assess the proportions of EPI and SEM. *Pdgfra* hypomorphic and null embryos had 45% and 43% of wild-type SEM content respectively (p<0.001, Fig. 3I). Deletion of one *Wt1* copy in *Pdgfra* null or hypomorphic embryos resulted in increased SEM content in about half of the embryos, but in the limited number of embryos that we were able to generate (n=6) this did not reach statistical significance (p=0.13, Fig. S7D). However, we also found that *Pdgfra^GFP/+^* heterozygotes, although grossly normal, also showed a reduction in SEM content to about 80% of controls (p=0.015, Fig. 4J). Deletion of one *Wt1* allele (*Pdgfra^GFP/+^ Wt1^GFPCre/+^*) was sufficient to restore the number of SEM cells to wild-type levels (p<0.0001, Fig. 4J).

Expression of epithelial and mesenchymal markers was assessed in EPI and SEM-like cells in *Wt1* mutants. E11.5 EPI cells (*Wt1*-GFP^+^ PODOPLANIN^high^ PDGFRB^−^) from *Wt1* nulls showed inappropriately high expression of ITGA4 and ALCAM, and low expression of PODOPLANIN (Fig. 5A-C). This marker pattern resembled that of wild-type PE (Fig. 5A-C), suggesting that differentiation of the EPI was impaired in *Wt1* mutants. E11.5 SEM-like cells in *Wt1* nulls (*Wt1*-GFP^+^ PODOPLANIN^low^ PDGFRB^+^) showed higher expression of PDGFRB (PE, p= 0.032; E10.5 EPI, p=0.0047) and lower expression of PODOPLANIN (PE, P= 0.0005; E10.5 EPI, p=0.0031) than in PE or the E10.5 EPI (Fig. 5C,D). This showed that the SEM-like cells had matured as mesenchymal cells. *Wt1* null SEM-like cells also showed a small but significant increase in apoptosis (Fig. 5E). The results of these experiments are consistent with a pro-epithelial role for WT1, since the absence of *Wt1* led to a less differentiated EPI but did not impair mesenchymal differentiation of SEM-like cells. The rescue of SEM cell numbers in Pdgfra heterozygotes also suggests that *Wt1* represses epicardial EMT.

**Figure 5.**
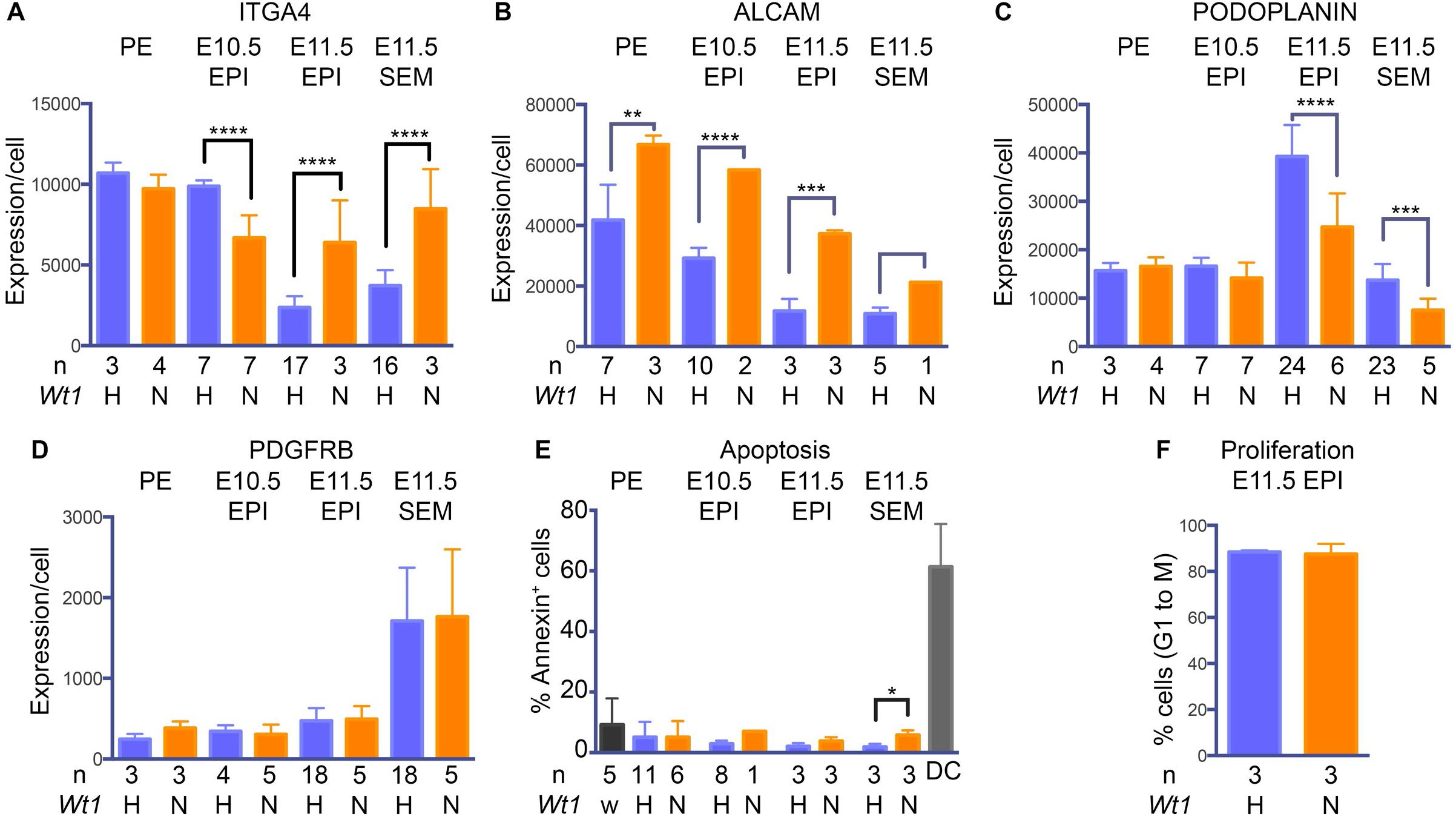
Early epicardial development is disrupted in *Wt1* null embryos. **A-D**. Average expression per cell (determined by flow cytometry) of ITGA4 (**A**), ALCAM (**B**), PODOPLANIN (**C**) and PDGFRB (**D**) during early epicardial development of *Wt1^GFPCre/+^* and *Wt1^GFPCre/GFPCre^* embryos. **E**. % annexin^+^ cells during early epicardial development of *Wt1^GFPCre/+^* and *Wt1^GFPCre/GFPCre^* embryos. DC= dead cells (fluorogold^+^). **F**. % Ki67^+^ cells within E11.5 EPI of *Wt1^GFPCre/+^* and *Wt1^GFPCre/GFPCre^* embryos. Genotypes: H: *Wt1^GFPCre/+^*; N: *Wt1^GFPCre/GFPCre^*. Mean ± SD are displayed. Statistics (Student’s t-test): * p<0.05, ** p <0.01, *** p <0.001, **** p<0.0001. n= number of independent samples.

### *Wt1* is required for transfer of proepicardial cells to the epicardium

The role of WT1 was also examined in the PE. There was no significant change in the total number of PE+EPI cells in *Wt1* nulls from E8.8 to E10.5 (Fig. 6A). Consistent with this, *Wt1* nulls showed no alteration in the frequency of apoptosis in PE and EPI cells (Fig. 5E) and proliferation of EPI cells was unaffected at E11.5 (Fig.5F). However, *Wt1-*GFP^+^ cell numbers increased in the *Wt1* null PE without increasing in the EPI, suggesting that WT1 is required for transfer from the PE to the heart (Fig. 6B,C). Of the PE markers we analyzed, only ALCAM was abnormal, showing increased expression in mutant PE cells (Fig. 5A-D). Expression of ITGA4, a predicted transcriptional target of WT1 (Kirschner et al., 2006), was unaffected in the *Wt1^GFPCre/GFPCre^* null PE (Fig. 5A) but reduced in the E10.5 EPI (Fig. 5A). A significant proportion of EPI cells lost all ITGA4 staining (Fig. 6D). Since *Itga4* is required for epicardial attachment to the heart (Sengbusch et al., 2002), this is likely to contribute to the low numbers of *Wt1*-GFP^+^ cells from E10.5 on (Fig. 6C,F) and the restoration of ITGA expression per EPI cell at E11.5 (Fig. 6A). However, reduced ITGA4 expression did not contribute to poor PE-to-EPI transfer in *Wt1* nulls because *Wt1-*GFP^+^ ITGA^high^ cells also accumulated in the PE (Fig. 6E,F). Despite having normal to excessive numbers of *Wt1-*GFP^+^ PE cells, *Wt1^GFPCre/GFPCre^* embryos showed a severe reduction in density and number of PE villi (n=4/4, Fig. 6G,H).

**Figure 6.**
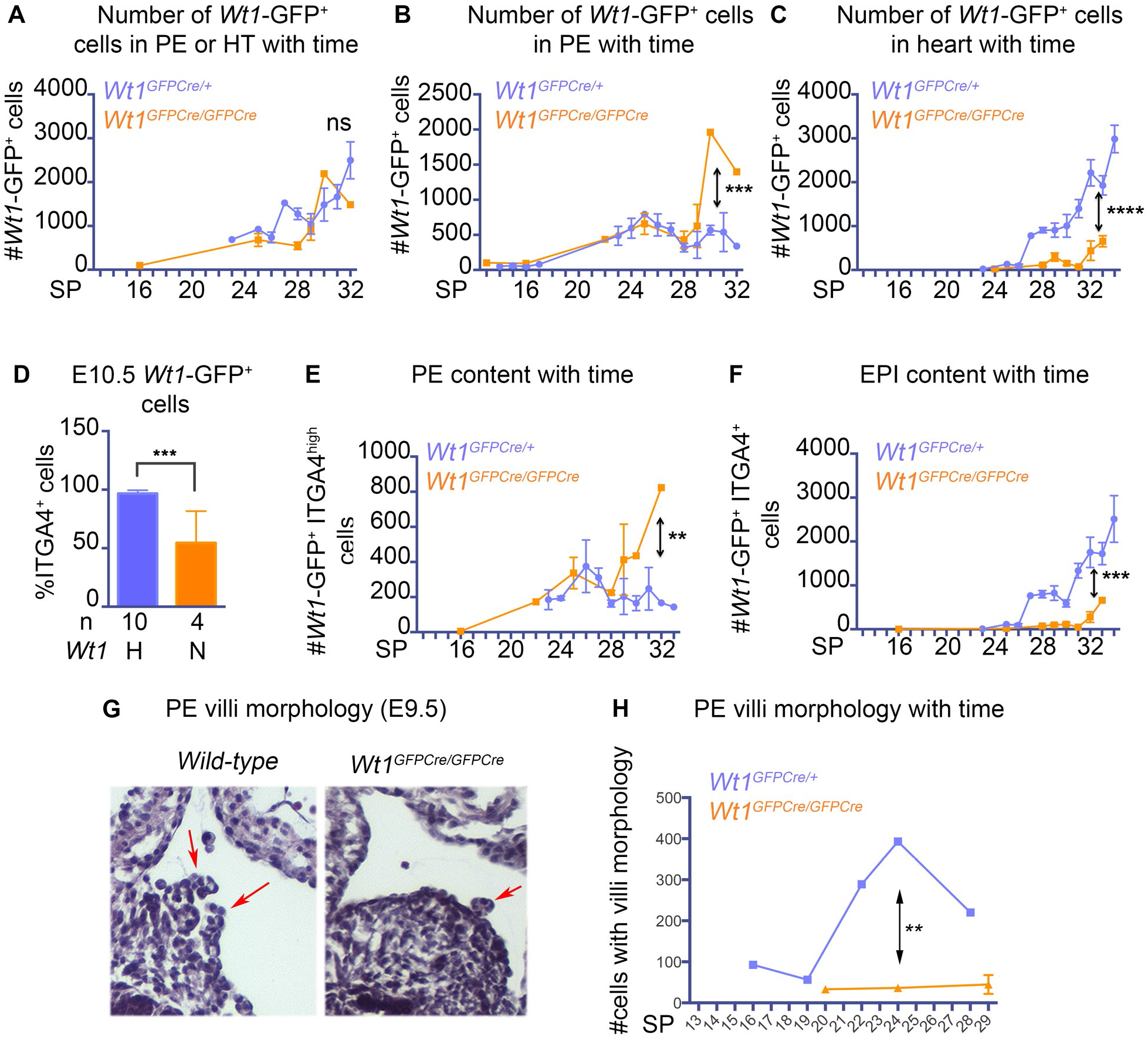
Proepicardial development is disrupted in *Wt1* null embryos. **A-C**. Total number of Wt1-GFP^+^ whether in PE or heart (**A**), PE (**B**) or heart (**C**) with time. **D**. % Wt1-GFP^+^ cells that express ITGA4 within the E10.5 heart. **E,F**. Number of proepicardial (ITGA4^high^) in PE (**E**) and epicardial cells (ITGA4^+^ ALCAM^+^) in heart (HT, **F**) in *Wt1^GFPCre/+^* and *Wt1^GFPCre/GFPCre^* embryos with time. SP = somite pairs. **G**. PE morphology on paraffin sections (Hematoxylin and Eosin counterstaining). Typical PE villi morphology is very rare in mutant embryos. **H**. Quantification of cells with villi morphology in *Wt1^GFPCre/+^* and *Wt1^GFPCre/GFPCre^* embryos with time. PE villi formation is absent and not delayed in mutants. Mean ± SD are displayed. Statistics: Student’s t-test: * p<0.05, ** p <0.01, *** p <0.001, **** p<0.0001.

## DISCUSSION

Epicardial development is a complex multistep process beginning with specification of the PE and culminating with differentiation of cardiac fibroblasts and smooth muscle cells. In order to understand the molecular mechanisms involved in these processes we developed methodology to identify, quantify and potentially isolate early epicardial derivatives throughout embryogenesis. We identified unique immunoprofiles which we used to perform a comprehensive and quantitative analysis of early epicardial development. This showed that PE formation, cell transfer to the EPI, formation of the SEM by EMT, and the accompanying marker expression are highly dynamic. In addition this method showed promise for quantifying other aspects of early cardiogenesis such as endothelium-to-mesenchyme transition (valve mesenchyme formation) and cardiac neural crest migration.

We tested this methodology on *Pdgfra* and *Wt1* mutants which are known to have epicardial phenotypes. Both genes are reported to be pro-mesenchymal and to promote EMT of the EPI (Martinez-Estrada et al., 2010; Smith et al., 2011; von Gise et al., 2011). We found supportive evidence for such a role for *Pdgfra* but not for *Wt1*. Embryos with less (hypomorphic and heterozygous) or no PDGFRA showed quantitative reductions in SEM. In contrast, we found that *Wt1* was required for formation of the PE villi, cell transfer to the heart, epithelial differentiation of the epicardium and epicardial EMT. *Wt1^−/−^* ventricles had a single external cell layer which has previously been interpreted as EPI without a SEM. Hence *Wt1*, like *Pdgfra*, was considered essential for epicardial EMT and interpreted as a pro-mesenchymal factor (Martinez-Estrada et al., 2010; von Gise et al., 2011). However, our quantitative analysis of *Wt1* null hearts showed that this external layer is composed of SEM-like mesenchymal cells and poorly epithelialized EPI cells. Defective formation of the SEM in *Pdgfra* heterozygous embryos was rescued when *Wt1* gene dosage was halved. Our experiments support previous in vitro reports suggesting that WT1 represses rather than promotes EMT (Bax et al., 2011; Takeichi et al., 2013).

*Wt1* nulls also had a severe reduction in PE villi, another an epithelial structure (Hirose et al., 2006; Manner, 1992; Nahirney et al., 2003). Collectively our data provides in vivo evidence for a pro-epithelial role for *Wt1* in the epicardial lineage. WT1 is known to be expressed in developing kidney cells transiting between mesenchymal and epithelial states, and in podocytes co-expressing mesenchymal and epithelial markers (Miller-Hodges and Hohenstein, 2012). *Wt1* is also required for the mesenchymal-to-epithelial transition (MET) in nephron formation (Davies et al., 2004). These observations suggest that *Wt1* might have a general pro-epithelial role in development.

EMT has been the focus of many studies as it is a mechanism widely used in developing embryos and disease. EMT is tightly regulated, with often some donor epithelium retained, while sufficient mesenchyme is produced. This is particularly true of epicardial EMT, which must spare epicardial integrity. Control of epicardial EMT reflects this duality, integrating the activity of epicardial maintenance factors such as WT1 with that of EMT inducers such as PDGF and TGF-beta pathways (von Gise and Pu, 2012) to generate coronary precursors.

## MATERIAL and METHODS

### Animal models

All transgenic (*Gata5Cre* (Zamora et al., 2007), *Tie2Cre* (Kisanuki et al., 2001), *Wnt1Cre* (Chai et al., 2000), Z/Red (Vintersten et al., 2004)) or knockin (*Nkx2-5^Cre^* (Stanley et al., 2002), *MesP1^Cre^* (Saga et al., 1999), *Wt1^GFPCre^* (Zhou et al., 2008), *Pdgfra*^*floxed*^ (Tallquist and Soriano, 2003), *R26R* (Soriano, 1999), *R26R-EYFP* (Srinivas et al., 2001)) lines had been maintained on a C57BL/6 background for at least 10 generations when obtained. They were kept on this background for the length of the study.

*Pdgfra*-H2BGFP mice (Hamilton et al., 2003) (referenced here as *Pdgfra^GFP^*), maintained on a C57BL/6 background, were bred once to FVB mice to generate F1 (C57BL/6 X FVB) females that were used for embryo collection. Survival and phenotype of null embryos was similar whether on C57BL/6 or a mixed C57BL/6 X FVB background, however the latter background was used for most analyzes due to a greater litter size.

All experiments were approved by the WEHI Animal Ethics Committee and conducted according to the Prevention of Cruelty to Animals Act 1986 (the Act) and the National Health and Medical Research Council Australian Code of Practice for the Care and Use of Animals for Scientific Purposes 8th edition (the NHMRC Code).

### Dissection

All embryos were scored for morphological criteria and health, and somite staged (when possible) under the microscope. Proepicardial regions were dissected and cleaned of sinus venosus and pericardium. Depending on the experiment, heart, out-flow tract or ventricles were dissected. When ventricle tissue was processed, endocardial cushions were removed to avoid contamination by valve tissues and neural crest derivatives.

### Flow cytometry

Dissociation protocols were optimized for each stage using survival of *Wt1*-GFP^+^ cells as a read-out. Dissected tissues were incubated in collagenase 2 (Worthington, 50u/ml) in PBS^−^ (without Ca and Mg), at 37 degrees for 10-30mins, depending on the stage/thickness of tissue. For late stage embryos or adult hearts, tissue was minced before enzymatic dissociation. PBS^−^ + 7%FCS was then added to the mix, followed by an optimized regime of gentle mechanical dissociation with a 1ml pipetman. Samples were washed, fitered and labelled for flow cytometry. Samples were incubated for 30mins to 1hr with primary or secondary antibodies in PBS^−^ + 7%FCS, followed by 2×2ml washes. Fluorogold (Sigma-Aldrich, 1/300) was used a viability marker. For intracellular FACS (Keratin labeling), samples were fixed 20mins in Cytofix/Cytoperm (BD Biosciences) on ice before staining. Primary antibodies were incubated overnight in wash buffer (BD Biosciences) at 4 degrees. Cells were then incubated with secondary antibodies for 4hrs on ice.

For proliferation studies, cells were fixed in ice cold 80% ethanol for 45mins and incubated with anti-Ki67 (BD Pharmingen kit) overnight. Cells were then stained with DAPI for 30mins at room temperature.

For all antibodies, pilot experiments included isotype controls. For Ki67 staining, the BD Pharmingen kit included an isotype control which was used in all experiments to ensure appropriate gating of Ki67^+^ cells.

All FACS amples were run on a Fortessa (BD Biosciences) and analyzed with FlowJO software.

### Statistics

Statistical analysis (unpaired Student’s t-test) was performed in Prism.

### Flow cytometry and immunofluorescence antibodies

Rat anti-mouse PDGFRA (eBiosciences, APA5), and PDGFRB (eBiosciences, APB5), rat anti-mouse ALCAM (eBiosciences, eBioALC48), hamster anti-mouse PODOPLANIN (eBiosciences, eBio8.1.1), rat anti-mouse ITGA4 (AbD Serotec, PS/2), rat anti-mouse PECAM (BD Pharmingen, MEC13.3), mouse anti-human Ki67 (BD Pharmingen), rabbit anti-GFP (Invitrogen, A11122), rabbit anti-mouse NKX2-5 (Santa Cruz sc-14033), rabbit anti-human WT1 (Santa Cruz, sc-192), rat anti-KRT8 (Troma I, DHSB), rat anti-KRT19 (Troma III, DHSB). Alexa conjugated secondary antibodies were from Invitrogen. Primary antibodies were used at 1/100 for FACS (1/400 for PODOPLANIN), and 1/50 for immunofluorescence. Specificity of staining was ascertained by isotype or secondary-only controls.

### Immunofluorescence

Samples were embedded in OCT following sucrose embedding, without prior fixation. 10μm sections were cut on a cryostat and stored at −80. After defrosting, sections were fixed for 3mins in 1% formalin, blocked for 1hr in PBS^−^ + 10% donkey serum and processed for antibody labeling. Antibodies incubation ranged from 1hr at room temperature to overnight at 4 degrees in PBS^−^ + 10% donkey serum. After antibody staining, slides were washed in PBS^−^ and incubated with DAPI (1/1000) before mounting in Immuno-Fluore (ImmunO). Samples were imaged on a LSM780 and processed with ImageJ or Imaris softwares.

### Whole-mount imaging

Hearts were dissected free from lung tissue and the outflow tract was sometimes removed to ensure proper mounting. Samples were fixed for 30mins in 2% paraformaldehyde 0.1% Tween 20 PBS^+^ (with Ca/Mg) at room temperature, then washed and blocked for 1hr in whole-mount buffer (WMB = PBS^−^ with 10%FCS + 0.6% Triton X-100). Overnight incubation with primary antibodies was followed by 6x 1hr washes in WMB, and incubation with secondary antibodies + DAPI when required. After washes, hearts were cleared overnight on glycerol gradient (Ferkowicz and Yoder, 2011), mounted between 2 microscope coverslips and imaged on a LSM780. Imaging data was processed with ImageJ or Imaris softwares.

### Histology

Embryos were fixed for 2 days in ice-cold 4% paraformaldehyde and processed for paraffin embedding. 7μm sections were counterstained with hematoxylin and eosin.

## Supporting information

SuppFig1

SuppFig2

SuppFig3

SuppFig4

SuppFig5

SuppFig6

SuppFig7

## List of Symbols and Abbreviations

EMT: Epithelial-to-Mesenchymal Transition
EPI: Epicardium
MET: Mesenchymal-to-Epithelial Transition
PE: ProEpicardium
PEP: ProEpicardial preparation
SEM: SubEpicardial Mesenchyme
VM: Valve Mesenchyme

## Acknowledgments

Assistance from Ferlene Ooi, Kath Colman, and advice from Kelly Rodgers, Anne Rios (imaging) and Tracey Baldwin (flow cytometry) was greatly appreciated. The authors are very grateful to Professor D. Hilton for freedom to operate and a great working environment.

## Author contributions

CB designed and performed experiments. OP designed experiments. CB and OP wrote the manuscript. BB, MM and LH provided technical assistance. ST and RPH read the manuscript.

## Supporting Information Legends

**Supp. Fig. 1. Quantitative analysis of early epicardial development**

**A-D**. Flow cytometry analysis of *Wt1*_*GFPCre/+*_ PE preparations (PEP). *Wt1*-GFP_+_ cells are PODOPLANIN_+_ PDGFRA_+_ ALCAM_+_ PDGFRB_−_. **E-H**. Flow cytometry analysis of *Wt1*_*GFPCre/+*_ E10.5 heart (HT). *Wt1*-GFP_+_ (EPI) cells are ALCAM_+_ (**E**) and the only cardiac cell type to be PODOPLANIN_+_ ITGA4+ or PECAM_−_ ITGA4_+_ (**F-H**). **I-L**. Flow cytometry analysis of *Wt1*_*GFPCre/+*_ E12.5 heart (HT). Gating on *Wt1*-GFP_+_ cells only. *Wt1*-GFP_+_ PDGFRA_+_ cells are PDGFRB_+_ (**I**) ITGA4_+_ (**J**). *Wt1*-GFP_+_ PODOPLANINhigh cells are *Wt1*-GFP_high_ (**K**) and ITGA4_low_ (**L**).

**Supp. Fig. 2. Relative EPI and SEM immunoprofiles using the *Pdgfra*-GFP knockin construct**

**A-F**. Expression profile of PDGFRA (**A**), KRT8 (**B**), KRT19 (**C**), PDGFRB (**D**), PODOPLANIN (**E**) and ITGA4 (**F**) with time in *Pdgfra*_*GFP/+*_ PE (ITGA4_high_) cells and ventricles preparations (E11.5-E13.5). Grey patterns: wild-type control re-GFP in PE. The boundary between GFP_+_ and GFP- cells in the heart is indicated in the first row of sample (**A**) (green line, based on wild-type samples). Green: Fluorescence minus one (FMO) negative control for KRT antibody stainings. Autofluorescent myocardium (MYO), *Pdgfra*-GFP_low_ EPI (blue arrow) and *Pdgfra*-GFP_high_ SEM (red gate) are indicated.

**Supp. Fig. 3. Identification of various cardiac lineages by flow cytometry**

**A-F**. Cells with a specific autofluorescence in the 488 (GFP) channel are ITGA4_−_ (**A**) PDGFRA_−_ (**B**), PECAM_−_ (**C**), *Nkx2-5*Cre-dsRed_+_ (**C**), VCAM_+_ (**D**), ALCAM_+_ (**E**) and PODOPLANIN_+_ (**F**) and therefore myocardial (MYO). **G-H**. Blood and endothelial derived lineages in *Tie2Cre*/+ *R26R-EYFP/+* hearts. Black boxes = *Tie2Cre*-YFP_mid_ PECAM_−_ Ter119_+_ deleted erythrocytes (E), grey box = *Tie2Cre*-YFP_−_ PECAM_−_ Ter119_+_ undeleted erythrocytes (UE), blue box = *Tie2Cre*-YFP_high_ PECAM_high_ Ter119_−_ endothelium, orange box = *Tie2Cre*-YFP_high_ PECAM_low_ Ter119_−_ valve mesenchyme. **I-J**. PECAM, ITGA4 and PDGFRA expression in *Tie2Cre*-YFP_high_ cells (gated of purple box in **H**). Blue = endothelium (PECAM_high_ ITGA4_low_ PDGFRA_−_); Orange = valve mesenchyme (PECAM_mid_ ITGA4_mid_, PDGFRAhigh); Green = rare EPI cells deleted by Tie2Cre (PECAM_−_ ITGA4_high_ PDGFRA_low_). **K**. *Wt1*-GFP expression in (PECAM_high_ ITGA4_low_ PDGFRA_−_, blue), (PECAM_mid_ ITGA4_mid_, PDGFRAhigh, orange) and EPI (ALCAM_+_ ITGA4_+_, red), confirming that both ENDO and VM populations as defined by those markers are *Wt1*-GFP_−_. **L**. ALCAM and ITGA4 expression in E10.5 *Tie2Cre*/+ *R26R-EYFP/+* hearts. Black = *Tie2Cre*-YFP_mid_ Ter119_+_ blood cells as in **G**; Blue, orange and green = *Tie2Cre*-YFP_high_ Ter119_−_ cells colour-coded as in **I**. The gate where epicardial cells normally can be found (EPI gate) is shown and devoid of blood or endothelial derivatives. **M**. LacZ (blue) and WT1 (brown) expression in a section from E9.5 *Tie2Cre/+ R26R/+* embryo, showing occasional recombination of PE cells with Tie2Cre: green arrow = LacZ+ WT1high PE cells; blue arrow = endocardial endothelium; black arrow = blood. **N**. PDGFRA, ITGA4, ALCAM and YFP expression in *Wnt1Cre/+ R26R-EYFP/+* hearts. Red = EPI cells (defined as ALCAM_+_ ITGA4_+_); Purple = neural crest-derived cells (*Wnt1*Cre-YFP_+_) which do not contaminated the EPI gate at that stage.

**Supp. Fig. 4. Quantification of valve mesenchyme and neural crest cells at in E10.5 hearts**.

**A**. Whole heart preparations of *Wnt1Cre/+ Rosa26YFP/+* embryos. YFP_+_ neural crest (NC) cells (blue) occupy the ENDO + VM gate. **B**. *Wnt1Cre/+ Rosa26-EYFP/+* hearts without OFT lack OFT cushions (by design) and all NC (blue) cells. **C**. Quantification of atrioventricular (AV) valve mesenchyme (AV VM) in hearts without OFT with time. As that of the epicardium (EPI), accumulation of AV VM is dynamic. SP: somite pairs number. **D**. Quantification of outflow tract (OFT) valve mesenchyme in *Wnt1Cre/+ Rosa26-EYFP/+* embryos. Mesenchymal cells (PDGFRA_+_ ITGA4_+_) are gated out of the ENDO + VM gate and separated into OFT VM (YFP_−_) and neural crest (YFP_+_).

**Supp. Fig. 5. Identification of SEM at E11.5 requires ventricle-only preparations in order to remove valve mesenchyme and neural crest-derived cells**.

**A,F**. ALCAM and ITGA4 expression in E11.5 wild-type whole hearts (**A**) and ventricle only preparations (**F**). EPI gate (red circle) is indicated. Note that the EPI gate is much cleaner in ventricle fractions. **B,C,G,H**. PECAM, ALCAM, ITGA4 and YFP expression in *Tie2Cre/+ R26R-EYFP/+* whole hearts (**B,C**) and ventricle-only preparations (**G,H**). Blue = endothelium; Orange = valve mesenchyme (VM); Green = rare *Tie2Cre*+ EPI cells (see Fig. S3M). EPI gate is indicated. Ventricle fractions have negligible VM content. **D,E**. Flow cytometry on *Wnt1Cre/+ R26R-EYFP/+* hearts. *Wnt1*Cre-YFP_high_ PDGFRA_+_ neural crest cells (purple) significantly contaminate the EPI gate at E11.5. **I,J**. Flow cytometry on ventricles or “everything else” (atria, atrioventricular groove [AVG], outflow tract [OFT]) preparations from *Wnt1Cre/+ R26R-EYFP/+* hearts showing that ventricle fractions are devoid of NC cells. **K-M** *Wt1*-GFP_+_ (EPI, red) and myocardial (MYO, teal) cells can be separated on a fluorogold vs GFP scatterplot. **N,O**. Flow cytometry on *Wt1*_*GFPCre/+*_ E11.5 ventricles. *Wt1*-GFP_+_ (EPI, red) and myocardial (MYO, teal) populations show significantly overlap, making a gating similar to Fig. 2B impossible with GFP_+_ cells.

**Supp. Fig. 6. *Gata5Cre/+ Pdgfra*_*floxed/floxed*_ embryos have no SEM and a stunted coronary network**.

**A-C**. PDGFRA and WT1 expression by immunofluorescence on sections in wild-type PE (**A**, E9.5), EPI (**B**, E10.5) and SEM (**C**, E12.5). **D**. Mean expression/cell for GFP epifluorescence, PODOPLANIN (PODO), ITGA4 and PDGFRB by flow cytometry in EPI of wild-types (w), *Pdgfra*_*GFP/+*_ (H) and *Pdgfra*_*GFP/GFP*_ (N) at E11.5. Mean ± SD are displayed. Statistics: Student’s t-test: no significant difference. **E,F**. Immunofluorescence on E13.5 sections. Genotypes are indicated. PECAM (green) highlights both the internal endocardial lining of the heart (EC), and the peripheral coronary endothelium (CE). NKX2-5 (pink) stains the myocardium. Epicardial lineages (EPI and SEM) are unstained in those conditions. Note the severe reduction in subepicardial mesenchyme and coronary endothelium in the mutant. **G,H**. Immunofluorescence on sections of E13.5 hearts. PECAM staining shows normal endocardial lining (EC) while the developing coronary network is much less dense and does not extend in mutants as far as in matched controls (red arrows). Note the absence of coronary endothelium at the level of the interventricular groove (IVG) in mutants. (yellow arrow).

**Supp. Fig. 7. *Wt1*_*GFPCre/+*_ embryos have normal early epicardial development**

**A**. Number of PE cells (ITGA4_high_) by flow cytometry in wild-type (grey) and *Wt1*_*GFPCre/+*_ embryos (blue). **B**. Number of EPI cells (PODOPLANIN_+_ ITGA4_+_) by flow cytometry in wild-type (grey) and *Wt1*_*GFPCre/+*_ embryos (blue). **C**. %SEM within the epicardial lineage at E11.5 by flow cytometry in wild-type (w, grey) and *Wt1*_*GFPCre/+*_ embryos (H, blue). Mean ± SD are displayed. Statistics: Student’s t-test: no significant difference. **D**. Proportion of SEM or SEM-like cells within the epicardial lineage of the ventricles at E11.5 (44-50sp) and E12.5 (51-57sp). Single embryos are represented. Statistics: Student’s t-test: no significant difference. Alleles for each gene: w = wild-type, H = heterozygote (either *Pdgfra*_*GFP/+*_ or *Wt1*_*GFPCre/+*_), Hp = hypomorph (*Pdgfra*_*GFP/floxed*_), N = null (either *Pdgfra*_*GFP/GFP*_ or *Wt1*_*GFPCre/GFPCre*_). **A,B**, each dot represents an embryo. **C,D**, n= number of independent samples.

## REFERENCES

Andrae, J., Gallini, R. and Betsholtz, C. (2008). Role of platelet-derived growth factors in physiology and medicine. Genes & development22, 1276–1312.

Asahina, K., Zhou, B., Pu, W. T. and Tsukamoto, H. (2011). Septum transversum-derived mesothelium gives rise to hepatic stellate cells and perivascular mesenchymal cells in developing mouse liver. Hepatology53, 983–995.

Bax, N. A., van Oorschot, A. A., Maas, S., Braun, J., van Tuyn, J., de Vries, A. A., Groot, A. C. and Goumans, M. J. (2011). In vitro epithelial-to-mesenchymal transformation in human adult epicardial cells is regulated by TGFbeta-signaling and WT1. Basic research in cardiology106, 829–847.

Cai, C. L., Martin, J. C., Sun, Y., Cui, L., Wang, L., Ouyang, K., Yang, L., Bu, L., Liang, X., Zhang, X., et al. (2008). A myocardial lineage derives from Tbx18 epicardial cells. Nature454, 104–108.

Chai, Y., Jiang, X., Ito, Y., Bringas, P., Jr., Han, J., Rowitch, D. H., Soriano, P., McMahon, A. P. and Sucov, H. M. (2000). Fate of the mammalian cranial neural crest during tooth and mandibular morphogenesis. Development127, 1671–1679.

Chao, R., Gong, X., Wang, L., Wang, P. and Wang, Y. (2015). CD71(high) population represents primitive erythroblasts derived from mouse embryonic stem cells. Stem cell research14, 30–38.

Chong, J. J., Chandrakanthan, V., Xaymardan, M., Asli, N. S., Li, J., Ahmed, I., Heffernan, C., Menon, M. K., Scarlett, C. J., Rashidianfar, A., et al. (2011). Adult cardiac-resident MSC-like stem cells with a proepicardial origin. Cell stem cell9, 527–540.

Davies, J. A., Ladomery, M., Hohenstein, P., Michael, L., Shafe, A., Spraggon, L. and Hastie, N. (2004). Development of an siRNA-based method for repressing specific genes in renal organ culture and its use to show that the Wt1 tumour suppressor is required for nephron differentiation. Human molecular genetics13, 235–246.

Dettman, R. W., Denetclaw, W., Jr., Ordahl, C. P. and Bristow, J. (1998). Common epicardial origin of coronary vascular smooth muscle, perivascular fibroblasts, and intermyocardial fibroblasts in the avian heart. Developmental biology193, 169–181.

Elliott, D. A., Braam, S. R., Koutsis, K., Ng, E. S., Jenny, R., Lagerqvist, E. L., Biben, C., Hatzistavrou, T., Hirst, C. E., Yu, Q. C., et al. (2011). NKX2-5(eGFP/w) hESCs for isolation of human cardiac progenitors and cardiomyocytes. Nature methods8, 1037–1040.

Ferkowicz, M. J. and Yoder, M. C. (2011). Whole embryo imaging of hematopoietic cell emergence and migration. Methods Mol Biol750, 143–155.

Hamilton, T. G., Klinghoffer, R. A., Corrin, P. D. and Soriano, P. (2003). Evolutionary divergence of platelet-derived growth factor alpha receptor signaling mechanisms. Mol Cell Biol23, 4013–4025.

Hirose, T., Karasawa, M., Sugitani, Y., Fujisawa, M., Akimoto, K., Ohno, S. and Noda, T. (2006). PAR3 is essential for cyst-mediated epicardial development by establishing apical cortical domains. Development133, 1389–1398.

Kirschner, K. M., Wagner, N., Wagner, K. D., Wellmann, S. and Scholz, H. (2006). The Wilms tumor suppressor Wt1 promotes cell adhesion through transcriptional activation of the alpha4integrin gene. The Journal of biological chemistry281, 31930–31939.

Kisanuki, Y. Y., Hammer, R. E., Miyazaki, J., Williams, S. C., Richardson, J. A. and Yanagisawa, M. (2001). Tie2-Cre transgenic mice: a new model for endothelial cell-lineage analysis in vivo. Developmental biology230, 230–242.

Komiyama, M., Ito, K. and Shimada, Y. (1987). Origin and development of the epicardium in the mouse embryo. Anatomy and embryology176, 183–189.

Kwee, L., Baldwin, H. S., Shen, H. M., Stewart, C. L., Buck, C., Buck, C. A. and Labow, M. A. (1995). Defective development of the embryonic and extraembryonic circulatory systems in vascular cell adhesion molecule (VCAM-1) deficient mice. Development121, 489–503.

Mahtab, E. A., Wijffels, M. C., Van Den Akker, N. M., Hahurij, N. D., Lie-Venema, H., Wisse, L. J., Deruiter, M. C., Uhrin, P., Zaujec, J., Binder, B. R., et al. (2008). Cardiac malformations and myocardial abnormalities in podoplanin knockout mouse embryos: Correlation with abnormal epicardial development. Developmental dynamics : an official publication of the American Association of Anatomists237, 847–857.

Manner, J. (1992). The development of pericardial villi in the chick embryo. Anatomy and embryology186, 379–385.

Martinez-Estrada, O. M., Lettice, L. A., Essafi, A., Guadix, J. A., Slight, J., Velecela, V., Hall, E., Reichmann, J., Devenney, P. S., Hohenstein, P., et al. (2010). Wt1 is required for cardiovascular progenitor cell formation through transcriptional control of Snail and E-cadherin. Nature genetics42, 89–93.

Mikawa, T. and Gourdie, R. G. (1996). Pericardial mesoderm generates a population of coronary smooth muscle cells migrating into the heart along with ingrowth of the epicardial organ. Developmental biology174, 221–232.

Miller-Hodges, E. and Hohenstein, P. (2012). WT1 in disease: shifting the epithelial-mesenchymal balance. The Journal of pathology226, 229–240.

Moore, A. W., McInnes, L., Kreidberg, J., Hastie, N. D. and Schedl, A. (1999). YAC complementation shows a requirement for Wt1 in the development of epicardium, adrenal gland and throughout nephrogenesis. Development126, 1845–1857.

Nahirney, P. C., Mikawa, T. and Fischman, D. A. (2003). Evidence for an extracellular matrix bridge guiding proepicardial cell migration to the myocardium of chick embryos. Developmental dynamics : an official publication of the American Association of Anatomists227, 511–523.

Plavicki, J. S., Hofsteen, P., Yue, M. S., Lanham, K. A., Peterson, R. E. and Heideman, W. (2014). Multiple modes of proepicardial cell migration require heartbeat. BMC developmental biology14, 18.

Rodgers, L. S., Lalani, S., Runyan, R. B. and Camenisch, T. D. (2008). Differential growth and multicellular villi direct proepicardial translocation to the developing mouse heart. Developmental dynamics : an official publication of the American Association of Anatomists237, 145–152.

Saga, Y., Miyagawa-Tomita, S., Takagi, A., Kitajima, S., Miyazaki, J. and Inoue, T. (1999). MesP1 is expressed in the heart precursor cells and required for the formation of a single heart tube. Development126, 3437–3447.

Sengbusch, J. K., He, W., Pinco, K. A. and Yang, J. T. (2002). Dual functions of [alpha]4[beta]1 integrin in epicardial development: initial migration and long-term attachment. J Cell Biol157, 873–882.

Singh, M. K., Lu, M. M., Massera, D. and Epstein, J. A. (2011). MicroRNA-processing enzyme Dicer is required in epicardium for coronary vasculature development. The Journal of biological chemistry286, 41036–41045.

Smart, N. and Riley, P. R. (2012). The epicardium as a candidate for heart regeneration. Future cardiology8, 53–69.

Smith, C. L., Baek, S. T., Sung, C. Y. and Tallquist, M. D. (2011). Epicardial-derived cell epithelial-to-mesenchymal transition and fate specification require PDGF receptor signaling. Circulation research108, e15–26.

Soriano, P. (1997). The PDGF alpha receptor is required for neural crest cell development and for normal patterning of the somites. Development124, 2691–2700.

Soriano, P. (1999). Generalized lacZ expression with the ROSA26 Cre reporter strain. Nature genetics21, 70–71.

Sridurongrit, S., Larsson, J., Schwartz, R., Ruiz-Lozano, P. and Kaartinen, V. (2008). Signaling via the Tgf-beta type I receptor Alk5 in heart development. Developmental biology322, 208–218.

Srinivas, S., Watanabe, T., Lin, C. S., William, C. M., Tanabe, Y., Jessell, T. M. and Costantini, F. (2001). Cre reporter strains produced by targeted insertion of EYFP and ECFP into the ROSA26 locus. BMC developmental biology1, 4.

Stanley, E. G., Biben, C., Elefanty, A., Barnett, L., Koentgen, F., Robb, L. and Harvey, R. P. (2002). Efficient Cre-mediated deletion in cardiac progenitor cells conferred by a 3’UTR-ires-Cre allele of the homeobox gene Nkx2-5. The International journal of developmental biology46, 431–439.

Takeichi, M., Nimura, K., Mori, M., Nakagami, H. and Kaneda, Y. (2013). The transcription factors Tbx18 and Wt1 control the epicardial epithelial-mesenchymal transition through bi-directional regulation of Slug in murine primary epicardial cells. PloS one8, e57829.

Tallquist, M. D. and Soriano, P. (2003). Cell autonomous requirement for PDGFRalpha in populations of cranial and cardiac neural crest cells. Development130, 507–518.

Vintersten, K., Monetti, C., Gertsenstein, M., Zhang, P., Laszlo, L., Biechele, S. and Nagy, A. (2004). Mouse in red: red fluorescent protein expression in mouse ES cells, embryos, and adult animals. Genesis40, 241–246.

Viragh, S. and Challice, C. E. (1981). The origin of the epicardium and the embryonic myocardial circulation in the mouse. The Anatomical record201, 157–168.

von Gise, A. and Pu, W. T. (2012). Endocardial and epicardial epithelial to mesenchymal transitions in heart development and disease. Circulation research110, 1628–1645.

von Gise, A., Zhou, B., Honor, L. B., Ma, Q., Petryk, A. and Pu, W. T. (2011). WT1 regulates epicardial epithelial to mesenchymal transition through beta-catenin and retinoic acid signaling pathways. Developmental biology356, 421–431.

Wang, J., Cao, J., Dickson, A. L. and Poss, K. D. (2015). Epicardial regeneration is guided by cardiac outflow tract and Hedgehog signalling. Nature522, 226–230.

Zamora, M., Manner, J. and Ruiz-Lozano, P. (2007). Epicardium-derived progenitor cells require beta-catenin for coronary artery formation. Proceedings of the National Academy of Sciences of the United States of America104, 18109–18114.

Zhou, B., Ma, Q., Rajagopal, S., Wu, S. M., Domian, I., Rivera-Feliciano, J., Jiang, D., von Gise, A., Ikeda, S., Chien, K. R., et al. (2008). Epicardial progenitors contribute to the cardiomyocyte lineage in the developing heart. Nature454, 109–113.

