## Supplementary figures and images for "Quantitation analysis by flow cytometry shows that *Wt1* is required for development of the proepicardium and epicardium"

### SuppFig1

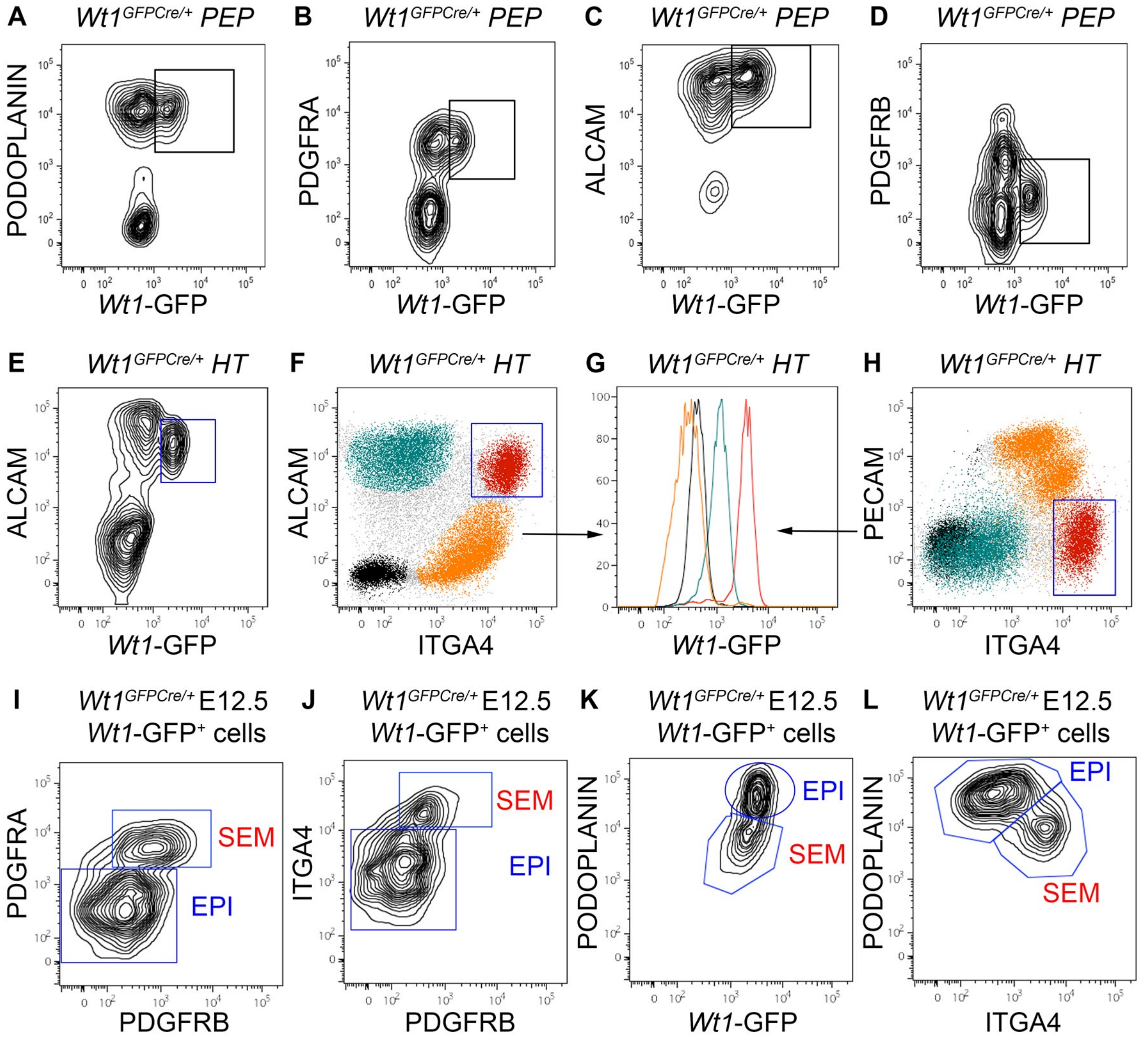


**Biben_Supplementary Figure 1**

### SuppFig2

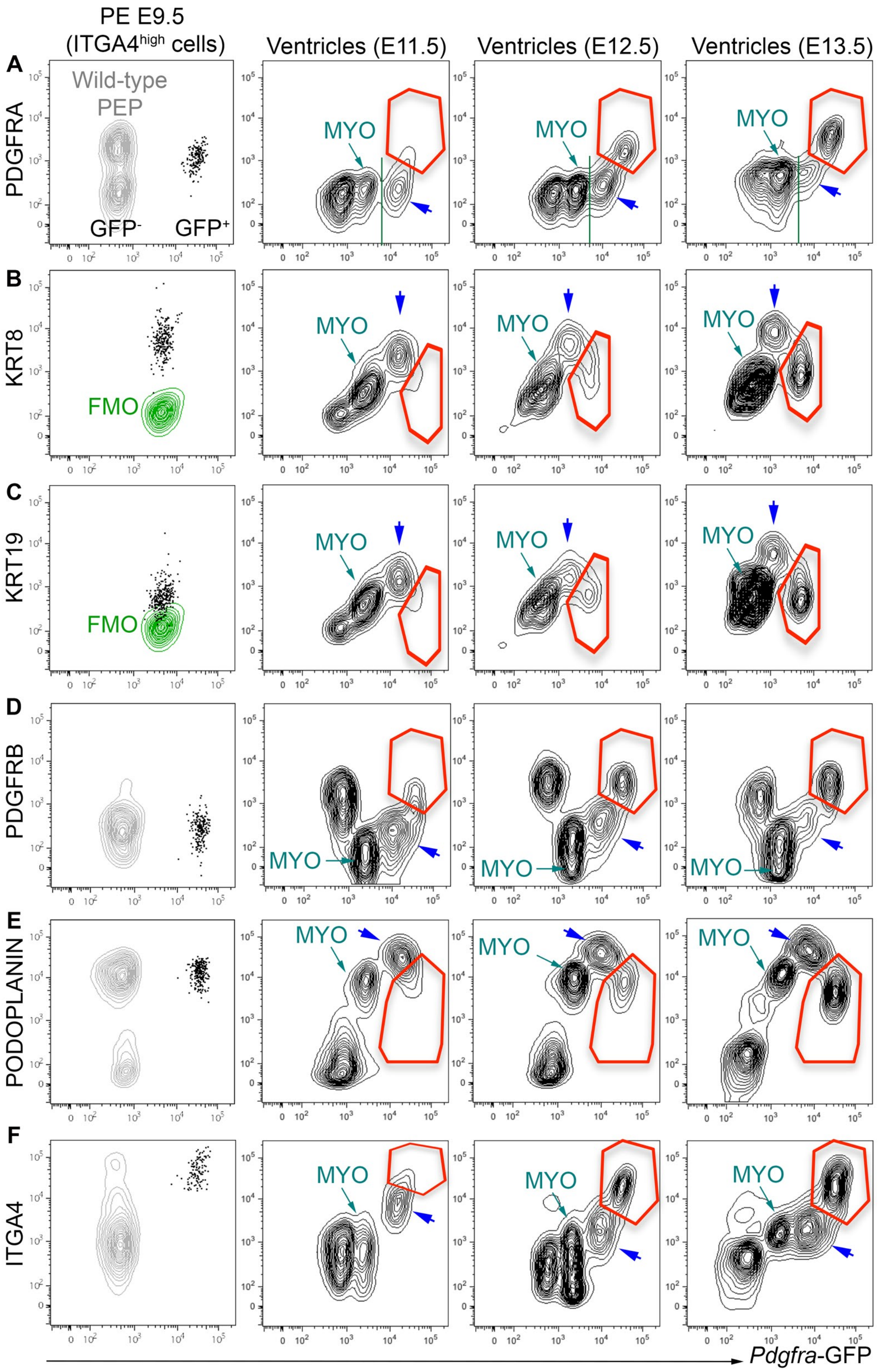


**Biben_Supplementary Figure 2**

### SuppFig3

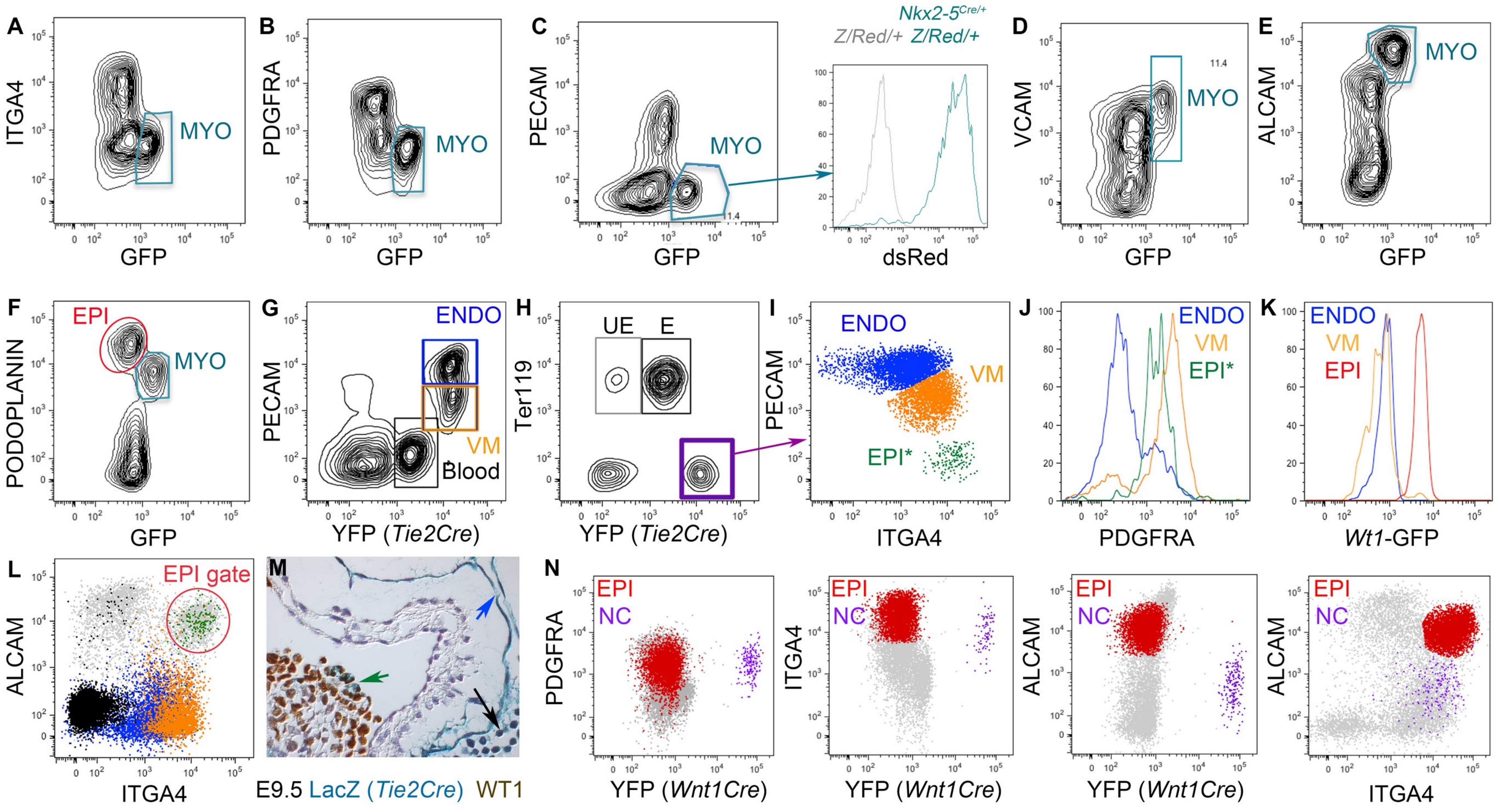


**Biben_Supplementary Figure 3**

### SuppFig4

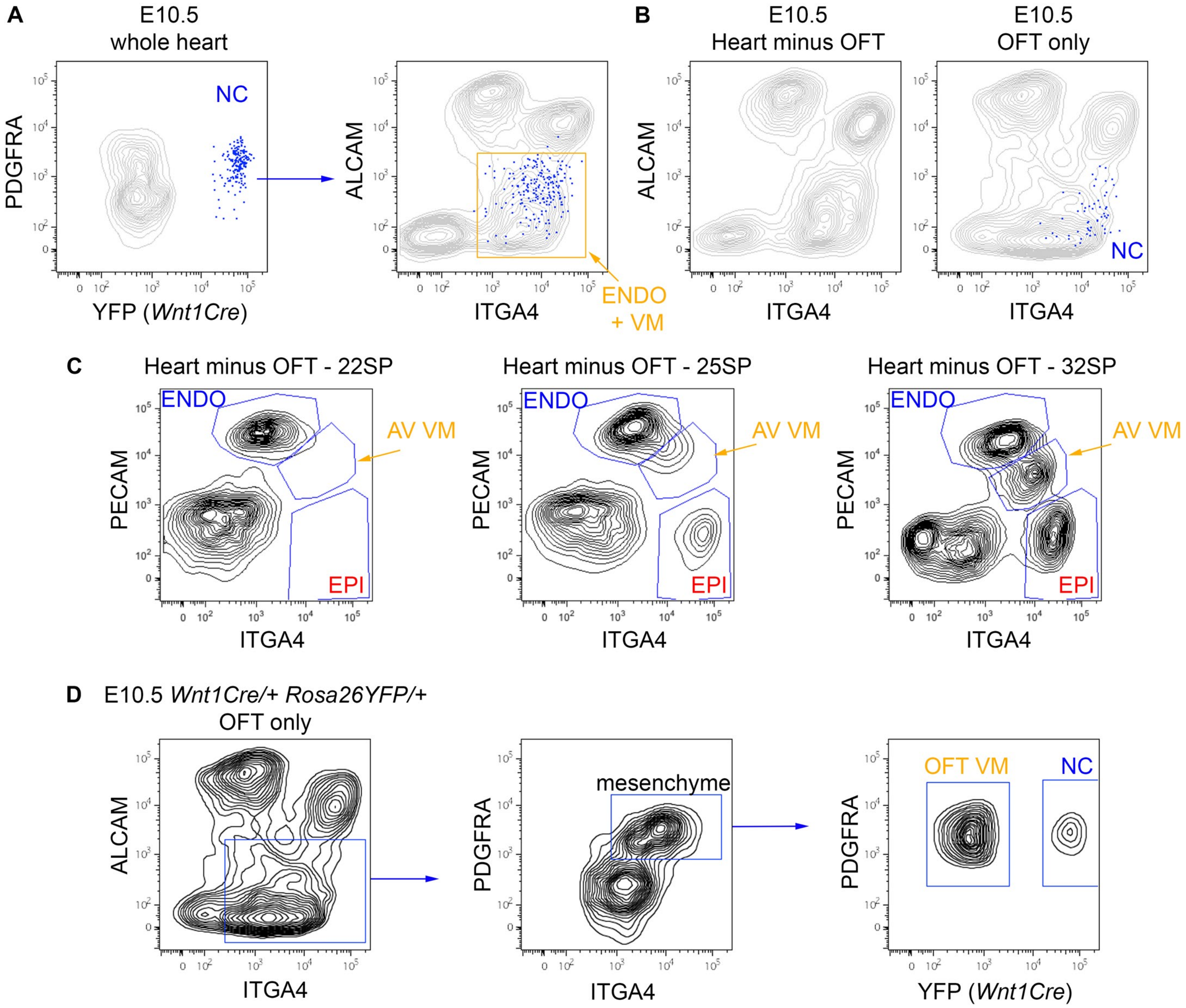


**Biben_Supplementary Figure 4**

### SuppFig5

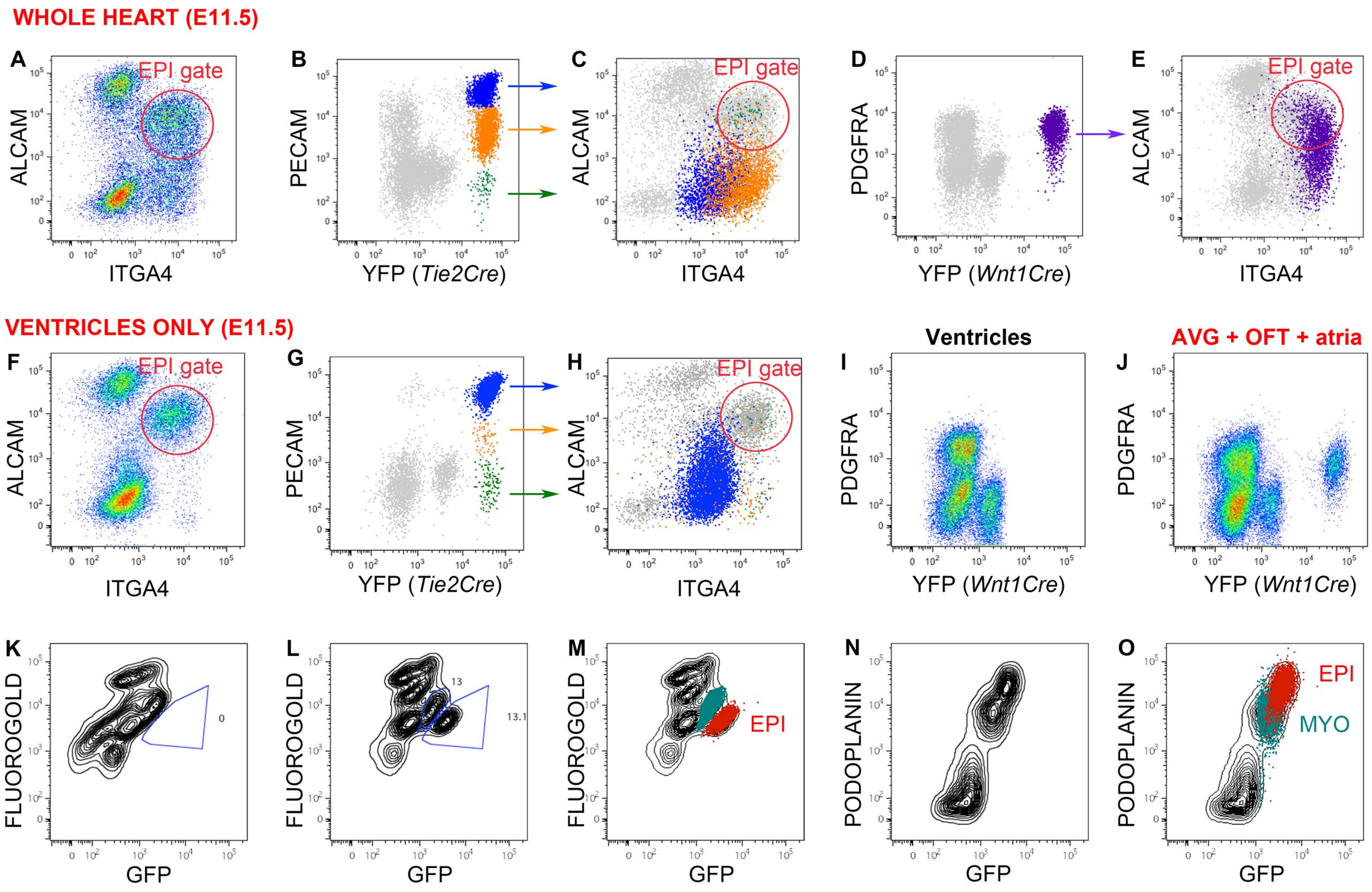


**Biben_Supplementary Figure 5**

### SuppFig6

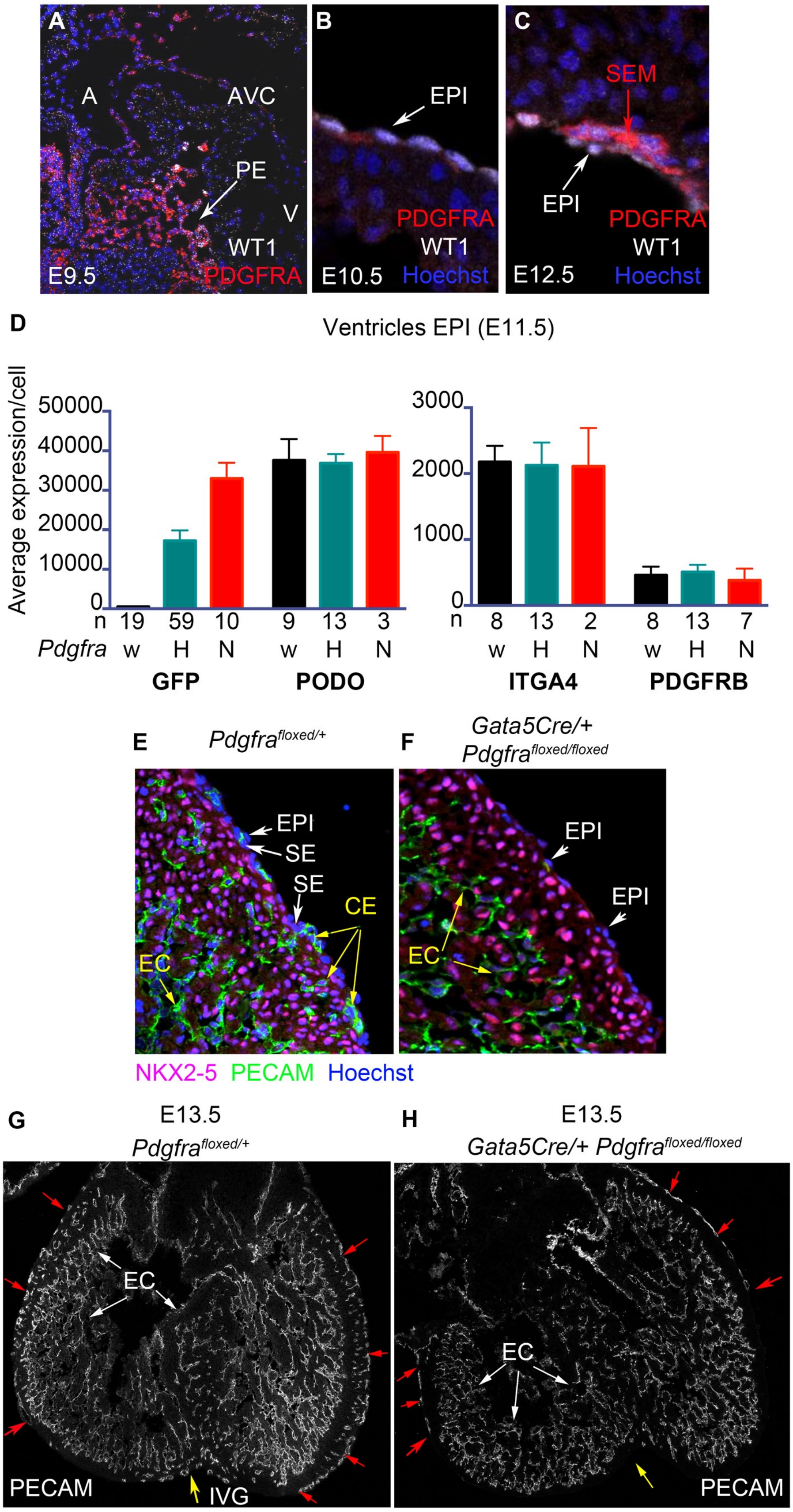


**Biben_Supplementary Figure 6**

### SuppFig7

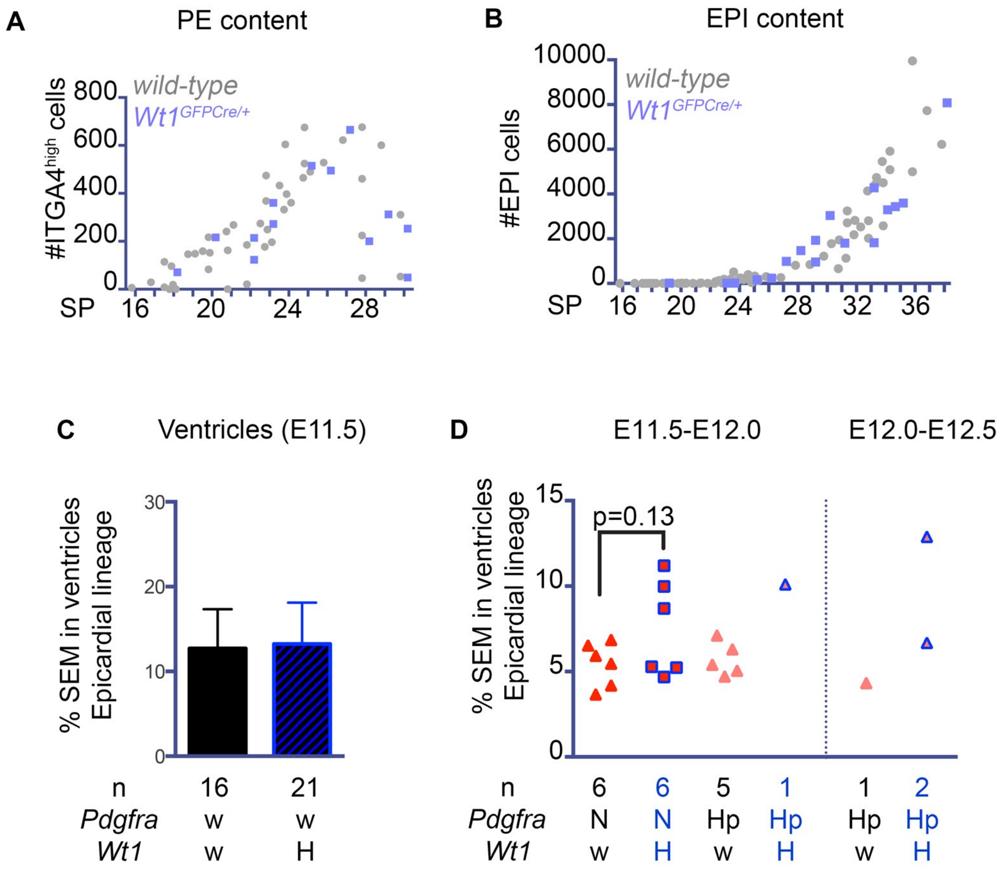


**Biben_Supplementary Figure 7**
